# Complex regulation of RETINOBLASTOMA-RELATED’s interactions with E2Fs via phosphorylation

**DOI:** 10.64898/2026.01.10.698770

**Authors:** Aladár Pettkó-Szandtner, Fruzsina Vadai-Nagy, Magdolna Gombos, Attila Hlacs, Eszter Molnár, Annamária Marton, Csaba Vizler, Rasik Shiekh Bin Hamid, Péter Kaló, Attila Fehér, Zoltán Magyar

## Abstract

Arabidopsis RETINOBLASTOMA-RELATED (RBR) regulates cell proliferation by interacting with E2F transcription factors and DIMERIZATION PARTNER, RB-LIKE, E2F AND MULTI-VULVAL CLASS B COMPLEX (DREAM) components. Although CDK-CYCD phosphorylation is believed to affect RBR’s E2F-binding capacity, the precise phosphorylation events inhibiting RBR’s cell cycle function remain unclear. This study found RBR phosphorylated at 13 of 16 CDK sites in Arabidopsis, with many phosphorylated forms still binding E2Fs. In contrast, multi-phosphorylated RBR forms with phosphorylated 911S site in *Arabidopsis thaliana* and corresponding sites in *Medicago truncatula* or *Brassica napus* do not co-purify with E2Fs and DREAM components but interact with RNA-binding proteins involved in post-transcriptional regulation through ribosomal biogenesis and protein translation. The 911S phosphorylation is high in proliferating cells and rapidly diminishes under DNA damage conditions, indicating its role in switching from proliferation to quiescence under stress. However, molecular modelling implies that this site is not accessible for phosphorylation if RBR is in complex with E2Fs. These findings suggest that different phosphorylation events inhibit RBR’s capacity to form complexes with E2Fs and to release E2Fs from RBR inhibition. We posit that multi-site phosphorylation coupled to 911S impedes free RBR’s binding to E2Fs and DREAM components, but this is not the initial inhibitory phosphorylation contributing to the disruption of RBR-E2F-DP complexes. Rather, it facilitates RBR interaction with proteins involved in post-transcriptional cell cycle regulation.

## Introduction

Plant growth and development depend on a balance between cell proliferation and differentiation, influenced by developmental and environmental signals. The plant homolog of the animal tumour suppressor Retinoblastoma (RB) plays a crucial role in the associated regulatory mechanisms (Harashima and Sugimoto, 2016). Plant RETINOBLASTOMA-RELATED (RBR) proteins, like their animal counterparts, regulate cell proliferation, quiescence, and survival (Desvoyes & Gutierrez, 2020). RBR proteins are structurally related to their animal relatives, consisting of conserved domains: the N-terminal domain, the central pocket domain (A and B), and the C-terminal domain. Both animal RB and plant RBR proteins bind E2F transcription factors to repress cell cycle entry (Desvoyes & Gutierrez, 2020; Zamora-Zaragoza et al., 2024). The pocket and C-terminal domains of animal RB associate with the transactivation domain and the so-called “marked box” region of E2Fs, stabilizing the E2F-RB complex (Dick & Rubin, 2013; Konagaya et al., 2024). Recent research shows that the canonical E2Fs (E2FA-C) in Arabidopsis are the main effectors of RBR in establishing transient and permanent cellular quiescence (Gombos et al., 2023). Further research has revealed that both animal RB and plant RBR proteins are involved in large multiprotein complexes, referred to as DIMERIZATION PARTNER, RB-LIKE, E2F AND MULTI-VULVAL CLASS B COMPLEX (DREAM – Sadasivam & DeCaprio, 2013; Kobayashi et al., 2015; Ning et al., 2020; Lang et al., 2021). It has been proposed that the plant DREAM-like complexes may form in response to DNA-damage and function as repressors, similar to their role in animals (Hafner et al., 2019; Lang et al., 2021). Therefore, it is well supported that RBR is a major growth regulatory protein in plants. Its activity is believed to be regulated by phosphorylation as previously demonstrated for animal RBs (Rubin et al., 2005; Rubin, 2013; Burke et al., 2014). Accordingly, there are 16 putative CYCLIN-DEPENDENT KINASE (CDK) phosphorylation sites at similar positions in the Arabidopsis RBR and animal RB proteins. However, only three RBR phospho-sites have been experimentally verified thus far (Reiland et al., 2009; Ábrahám et al., 2011; Willems et al., 2020; Desvoyes & Gutierrez, 2020; Zamora-Zaragoza et al., 2024, and Fig. 1). One of these confirmed phospho-sites is the 911 serine residue (911S) of the Arabidopsis RBR, and its phosphorylation could be monitored by using a human phospho-RB specific antibody (recognizing the phosphorylated 807S/811S residues in the human RB – Ábrahám et al., 2011). The mutation of 911S of the Arabidopsis RBR to alanine has been shown to prevent the recognition of the protein by this specific antibody (Wang et al., 2014). This site was found to be highly phosphorylated in an Arabidopsis line overexpressing CYCD3;1, while its level dropped in another line ectopically expressing the CDK-inhibitor KIP-RELATED PROTEIN 2 (KRP2) supporting that its phosphorylation is regulated by a CDKA-CYCD-KRP module (Magyar et al., 2012). Additionally, RBR was shown to be highly phosphorylated at this site in proliferating cells of developing young leaves (Őszi et al., 2020). Within synchronized cycling cells of a *Medicago sativa* suspension culture, the P-911S phosphorylation signal was detected in cells advancing through the S-phase reaching its maximum during the G2 and M phases (Ábrahám et al., 2011). These data collectively indicate that RBR is phosphorylated at this site in dividing plant cells, and it remains phosphorylated throughout the cell cycle. Moreover, immunprecipitated E2FA and E2FB proteins could not pull down the P-911S RBR protein from developing young leaves (Magyar et al., 2012; Őszi et al., 2020). However, it is unknown if it was the direct consequence of 911S phosphorylation and/or the simultaneous phosphorylation of RBR at other sites. Recent research indicates that combinatorial phosphorylation of Arabidopsis RBR sites distinctly modulates its roles in cell proliferation, stem cell fate, and cell death regulation (Zamora-Zaragoza et al., 2024).

**Figure 1.**
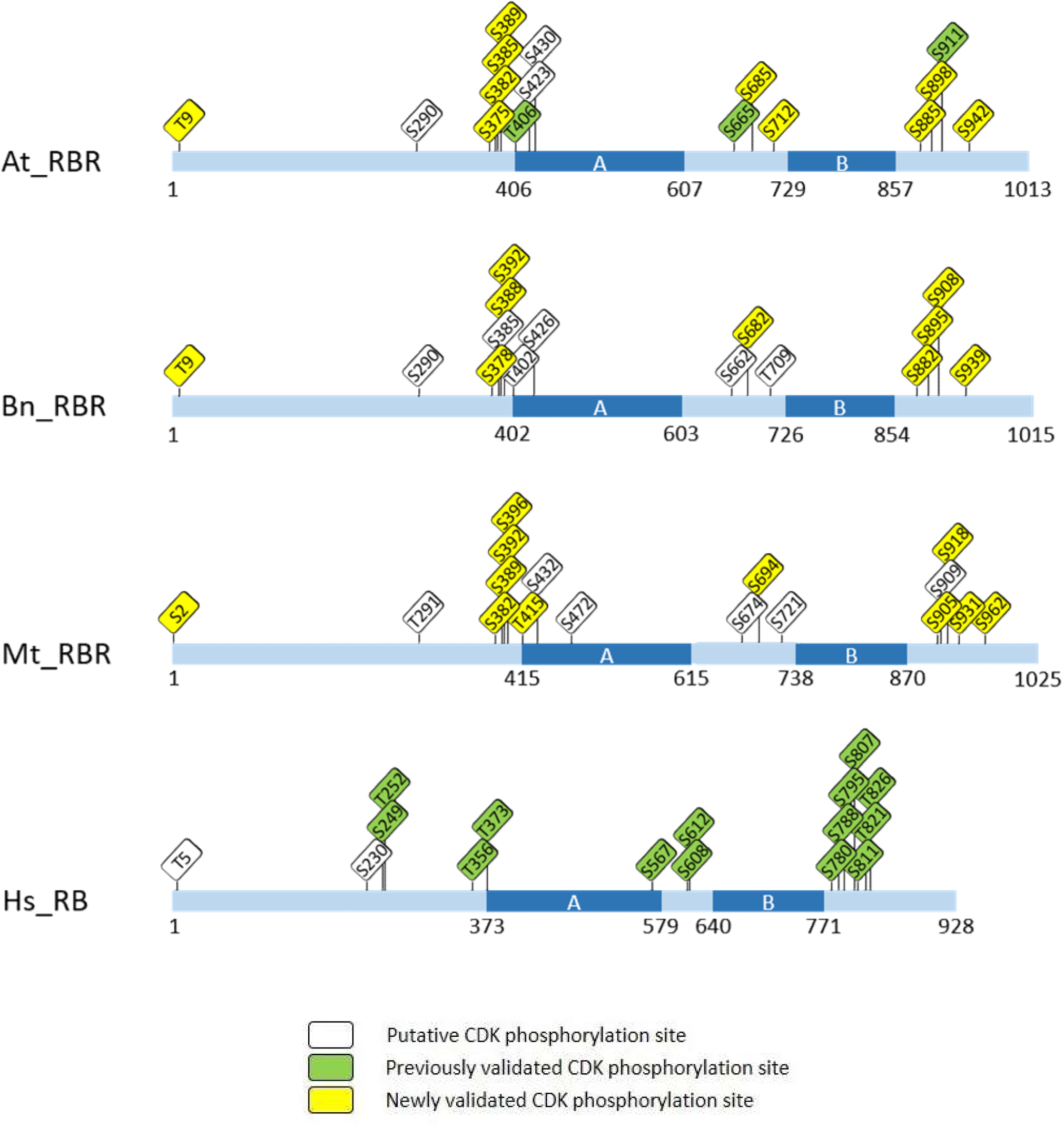
*In silico* predicted and experimentally validated CDK phosphorylation sites in plant RBR proteins in comparison to the mammalian RB. Domain organization and phosphorylation site distribution in *Arabidopsis thaliana* (At_RBR), *Brassica napus* (Bn_RBR) and *Medicago truncatula* (Mt_RBR) retinoblastoma-related proteins in comparison with those of the human RB (Hs_RB). For the sake of simplicity, we selected one of the six BnRBR proteins from Brassica specifically the BnaA01g30730D. The A and B (dark blue) domains define the “pocket” domain implicated in E2F/DP binding. The CDK/cyclin consensus phosphorylation sites (rectangles) are also indicated; the previously validated phosphorylation sites are shown in green, the sites newly validated by this study are in yellow, while the predicted but not yet validated phosphorylation sites are marked with white boxes.

Earlier animal studies showed that CDK phosphorylation of RB at multiple unique sites caused distinct structural changes, leading to E2F dissociation (Dick and Rubin, 2013; Rubin et al., 2020). However, recent findings have revealed that mono-phosphorylation of animal RB at any of the 14 unique CDK sites does not inhibit E2F binding (Narashima et al., 2014; Sanidas et al., 2019). Instead, these mono-phosphorylated forms (mP-RB) not only repress E2F target genes but also target specific gene sets with functions beyond the cell cycle. For example, mP-RB at S811 was shown to enhance mitochondrial activity and promote RB assembly with factors in the NuRD chromatin remodelling complex (Sanidas et al., 2019). Furthermore, a substantial portion of the RB interactome specifically binds to hyper-phosphorylated RB (Sanidas et al., 2019). In contrast to the mono-phosphorylated forms, the multi-phosphorylated RB protein loses its ability to form complexes with E2Fs and cannot participate in large multi-protein assemblies such as DREAM (Sadasivam & DeCaprio, 2013; Rubin et al., 2020; Zaragoza et al., 2021). However, it remains unclear whether this alters RB activity in association with other proteins involved in processes distinct from transcriptional regulation. The structural similarity between animal RB and plant RBR proteins suggests similar effects on plant RBR upon phosphorylation at conserved sites (Fig. 1). However, the extent to which plant RBR proteins function similarly to animal RBs remains unexplored.

This investigation shows that plant RBR proteins undergo phosphorylation at many putative CDK sites in planta, without inhibiting their interaction with E2F TFs. Purification of RBR from Arabidopsis seedlings using a phospho-specific antibody targeting the conserved P-911S site revealed an enriched pool of phosphorylated RBR forms phosphorylated also at several other sites. Interestingly, these P-911S-coupled multiP-RBR forms could not interact with E2Fs or other DREAM components in Arabidopsis seedlings, *Medicago truncatula* root tips, or proliferating *Brassica napus* leaf cells. However, they were found to be associated with proteins that possess RNA-binding activity and are involved in post-transcriptional regulation through ribosomal biogenesis and protein translation suggesting a phosphorylation-dependent switch in RBR’s function. We propose that multi-site phosphorylation in conjunction with 911S cannot be the initial step in disrupting RBR-E2F-DP complexes as the 911S site remains inaccessible to CDKs when the RBR is in complex with E2F-DPs. These findings suggest that different phosphorylation events might control RBR’s ability to form complexes with E2Fs or release them. Moreover, we found that RBR phosphorylation at 911S shows a rapid decrease under DNA damaging conditions, coinciding with growth arrest. This indicates the importance of RBR phosphorylation in controlling cell proliferation during stress.

## Results

### Validation of conserved phospho-sites in plant RBR proteins

In a previous study, we conducted a meta-analysis on a vast set of mass spectrometry data generated from co-immunoprecipitation experiments of proteins associated with the C-terminally GFP-tagged Arabidopsis RBR under diverse growth-promoting and growth-restricting environmental conditions (Lang et al., 2021). To determine the phosphorylation sites in the RBR protein, we reanalysed these immunoprecipitates (IP-s) derived from young seedlings. Sixteen phosphorylated sites were identified in RBR; most of these sites were putative CDK target sites (Fig. 1), and three among them have previously been verified (406T, 665S, 911S – Desvoyes and Gutierrez 2020). Three *in silico* predicted CDK recognition sites (290S, 423S, 430S) were not confirmed by our analyses, while three of the phosphorylated sites were not in CDK motifs but were in their proximity (708S, 893S, 903S). Through immunoprecipitation of endogenous non-tagged RBR from seven-day-old Arabidopsis wild type seedlings, we further validated 11 phospho-sites in the endogenous RBR protein (Table 1). Utilizing the phospho-specific anti-P-Rb^807/811S^ antibody in both ectopic RBR-GFP expressing and wild-type Arabidopsis lines, all the 13 CDK sites were observed to be phosphorylated (Table 1, Fig. 1). In summary, we experimentally confirmed the phosphorylation of thirteen CDK motifs in the RBR proteins immunoprecipitated from Arabidopsis seedlings.

**Table 1.**
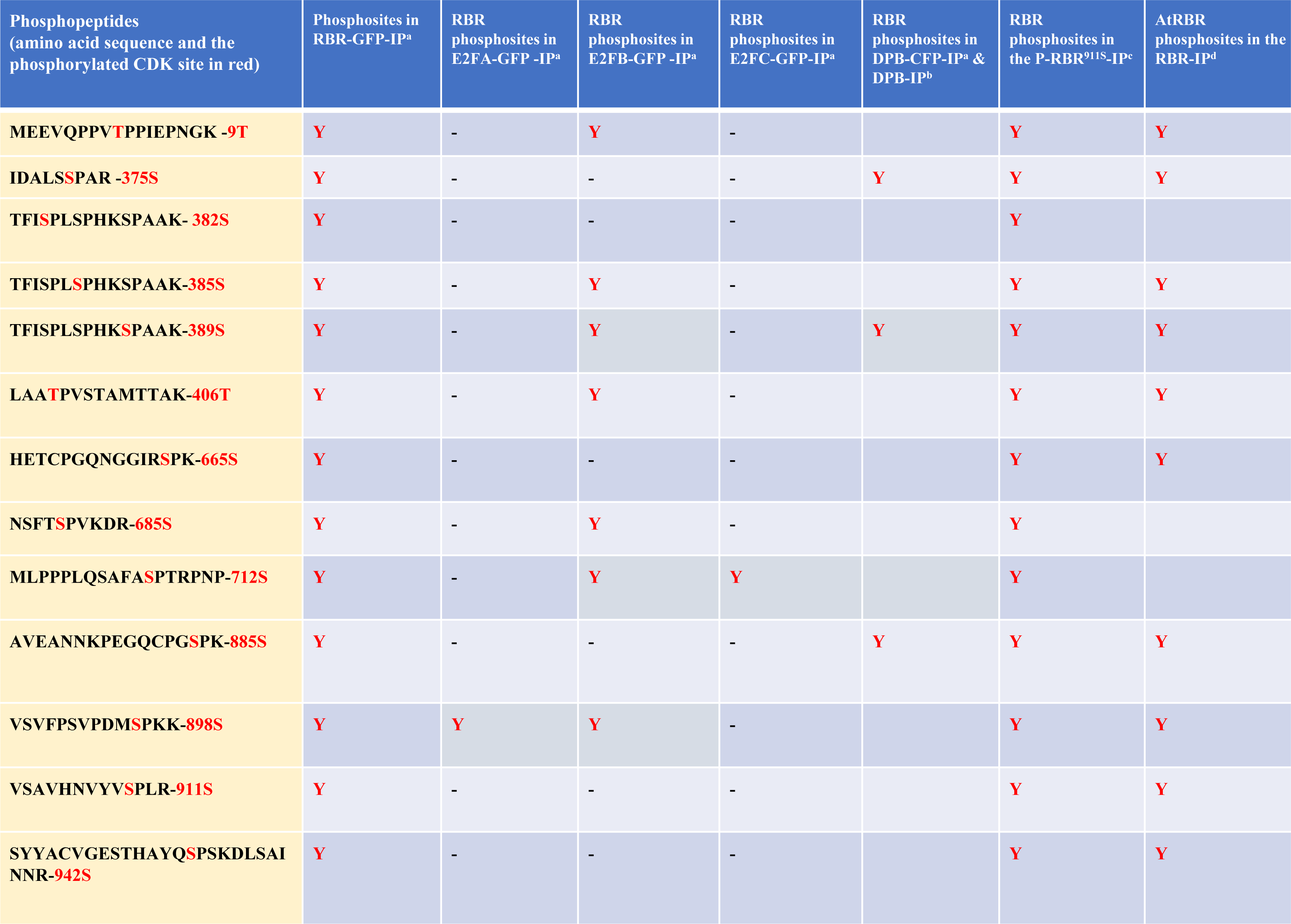
Phosphorylated CDK sites identified by mass spectrometry in AtRBR protein. Phosphorylated CDK sites were identified in RBR-GFP or RBR proteins immunoprecipitated (IP) using anti-GFP^a^, anti-AtRBR^d^, or anti-P-RBR^c^ (Anti-human P-RB^807/811S^) antibodies or in co-immunoprecipitated complex with E2FA/B/C-GFP or DPB-GFP/DPB using anti-GFP or anti-DPB^b^ antibodies.

To confirm the Arabidopsis data, RBR proteins were immunprecipitated from *M. truncatula* roots tips and *B. napus* young leaves using the anti-P-Rb^807/811S^ antibody (Table 2) and determined the phosphorylation sites through subsequent MS analyses. MtRBR possesses 17 putative CDK sites at similar positions than the Arabidopsis RBR and 11 of those were found to be phosphorylated in the immunprecipitate (Fig. 1; Suppl. Table 1). Due to the allopoliploid nature of *B. napus*, six BnRBR-related proteins were efficiently precipitated from the leaf samples of this species (Table 2; Suppl. Data 1). These six different but strongly homologous proteins were found to be identical to the predicted BnRBR proteins originating from their genomic clones, indicating that they exhibit similar accumulation patterns in young Brassica leaves. Putative CDK sites were located at similar positions in the Brassica RBR proteins as in the Arabidopsis, and Medicago ones. Nine of these CDK sites were found to be phosphorylated in the RBRs from the *B. napus* leaves (Fig. 1; Suppl. Table 1).

**Table 2.**
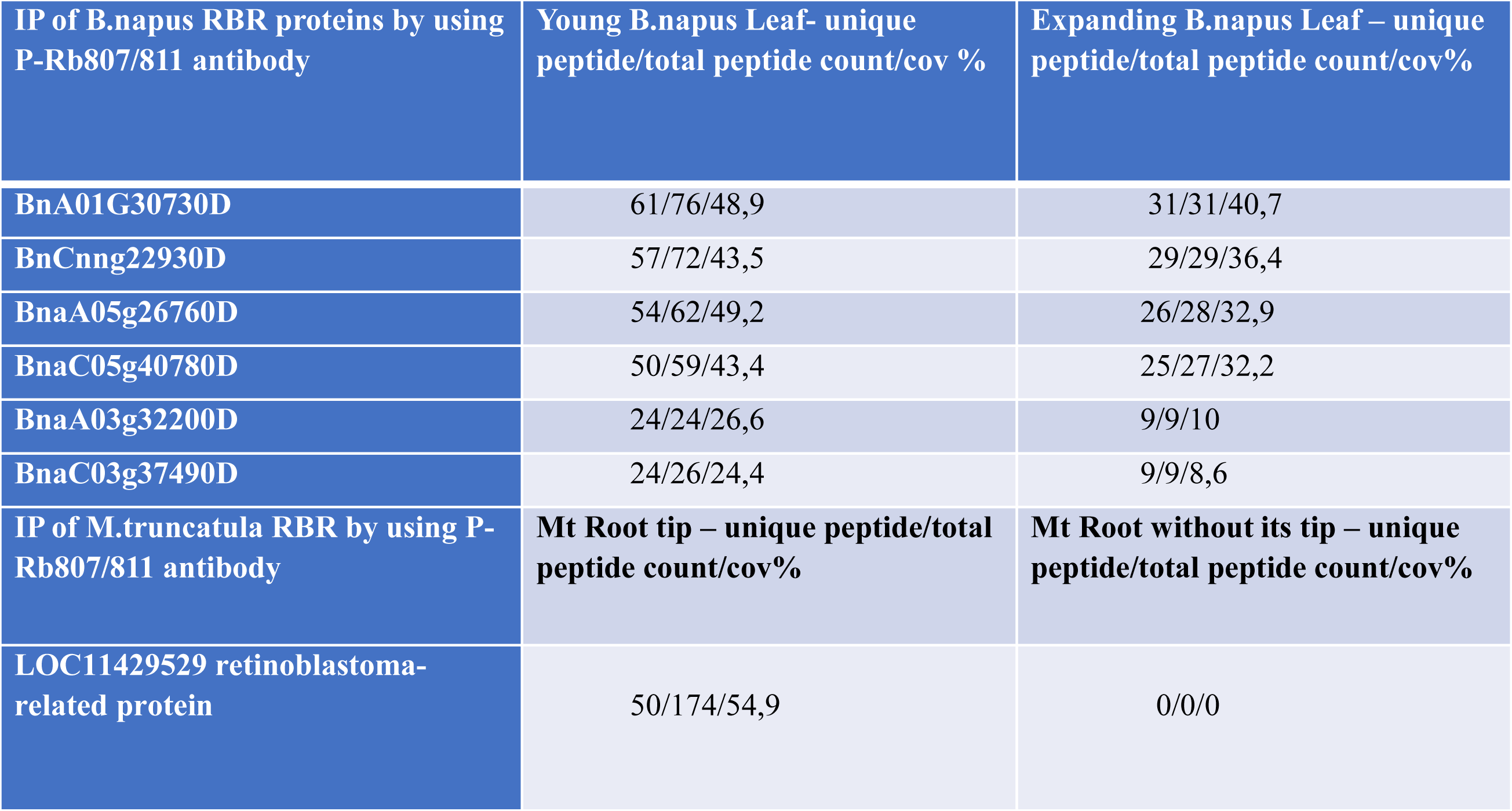
RBR proteins were immunoprecipitated from Medicago and Brassica species. RBR proteins were effectively purified from the proliferative first leaf of *Brassica napus* and root tip of *Medicago truncatula* utilizing a phospho-specific human Rb antibody (anti-pRb^807/811^). IP = immunprecipitated, cov =coverage

| **IP of B.napus RBR proteins by using P-Rb807/811 antibody** | **Young B.napus Leaf- unique peptide/total peptide count/cov %** | **Expanding B.napus Leaf – unique peptide/total peptide count/cov%** |
| --- | --- | --- |
| **BnA01G30730D** | 61/76/48,9 | 31/31/40,7 |
| **BnCnng22930D** | 57/72/43,5 | 29/29/36,4 |
| **BnaA05g26760D** | 54/62/49,2 | 26/28/32,9 |
| **BnaC05g40780D** | 50/59/43,4 | 25/27/32,2 |
| **BnaA03g32200D** | 24/24/26,6 | 9/9/10 |
| **BnaC03g37490D** | 24/26/24,4 | 9/9/8,6 |
| **IP of M.truncatula RBR by using P-Rb807/811 antibody** | **Mt Root tip – unique peptide/total peptide count/cov%** | **Mt Root without its tip – unique peptide/total peptide count/cov%** |
| **LOC11429529 retinoblastoma-related protein** | 50/174/54,9 | 0/0/0 |

This corroborates that plant RBR proteins, in analogy to their animal counterparts, are regulated by phosphorylation.

### Various phosphorylated forms of Arabidopsis RBR occur in complexes with E2F transcription factors

Previous research has demonstrated that GFP-tagged RBR exhibited binding affinity for E2F, DP, and the DREAM components (Lang et al., 2021). According to the current cell cycle regulation model, phosphorylation of RBR may inhibit its binding to E2F-DP heterodimers and DREAM components (Rubin et al., 2020; Zamora-Zaragoza et al., 2024). We conducted a reanalysis of the previously established interactome of E2FA, E2FB, E2FC, and DIMERIZATION PARTNER B (DPB) proteins to investigate whether E2Fs can form complexes with phosphorylated RBR. The dataset was derived from seedlings expressing the three distinct canonical E2Fs and DPB, each under the control of their respective promoters, tagged with GFP and CFP, respectively. The fusion proteins were immunoprecipitated using an anti-GFP antibody, along with their associated protein partners (Lang et al., 2021). Furthermore, immunoprecipitation was conducted in young Arabidopsis leaves utilizing an anti-DPB specific antibody to identify the associated partners (see later in Fig. 4C). These datasets were examined for the presence of phosphorylated RBR peptides. Seven phosphorylated CDK sites of RBR were identified in association with E2Fs and two additional phospho-sites with DPB. All seven phospho-peptides could be detected in the E2FB-GFP interactome, of which two distinct sites were detected in association with E2FA-GFP (898S) and E2FC-GFP (712S), respectively (Table 1). Additionally, three phospho-sites were associated with DPB, of which two (374S and 885S) were not observed in association with any E2F. This observation suggests that phosphorylation of RBR may differentially influence its binding affinity towards E2Fs with E2FB being the most favoured partner suggesting a specific functional relationship (Lang et al., 2021). Alternatively, the differences may reflect the abundance of E2F proteins in planta with E2FB being the most abundant and E2FC being the least abundant E2F protein (Wang et al., 2015). The finding also indicates that phosphorylation of RBR at various sites does not necessarily prevents its interaction with E2Fs. These phosphorylations likely contribute to the fine control of RBR’s activity.

### The phosphorylation of plant RBRs at sites equivalent to AtRBR’s 911S is connected to cell proliferation

A phospho-site-specific antibody against the animal P-807/811S sites has been proven to specifically recognize the corresponding phosphorylated site in *Arabidopsis thaliana* (911S) and *Medicago sativa* (931S) RBRs (Ábrahám et al., 2011; Magyar et al., 2012; Wang et al., 2014). Using this antibody, phosphorylation of RBR at the RBR^911S^ site has previously been found to be most prominent in proliferating organs and cells of Arabidopsis (Ábrahám et al., 2010; Magyar et al., 2012; Őszi et al., 2020; Gombos et al., 2023). To further support these observations, we examined the phosphorylation of AtRBR at the 911S site using western blot assay after treating 5-day-old seedlings with 50 µM cisplatin for 8 and 24 hours, inhibiting proliferative activity due to DNA damage response (De Schutter et al., 2007), and compared these samples to those of non-treated control seedlings (Fig. 2). As Figure 2A illustrates, root growth was significantly impeded in the presence of cisplatin and ceased entirely 48 hours after the application. Previous research has demonstrated that RBR specifically forms complexes with DREAM components, including E2FB, when seedlings are exposed to this DNA damaging agent, suggesting that RBR activity increases due to reduced phosphorylation (Lang et al., 2021). In contrast to the control non-treated sample where the P-RBR^911S^ signal increased at 8 hours, the level of P-RBR^911S^ declined in the presence of the drug and was almost completely diminished after 24 hours of treatment (Fig. 2B-D). Unlike the P-RBR^911S^ forms, both RBR and E2FB proteins exhibited relatively constant levels in this experiment (Fig. 3). In conclusion, these experiments provide additional proof that phosphorylation of RBR at the 911S site strongly and positively correlates with cell proliferation. The result agrees with the model that the suppression of CDK kinases and activation of phosphatases results in fully active, non-phosphorylated RBR during the DNA damage response, inhibiting proliferative activity (De Schutter et al., 2007).

**Figure 2.**
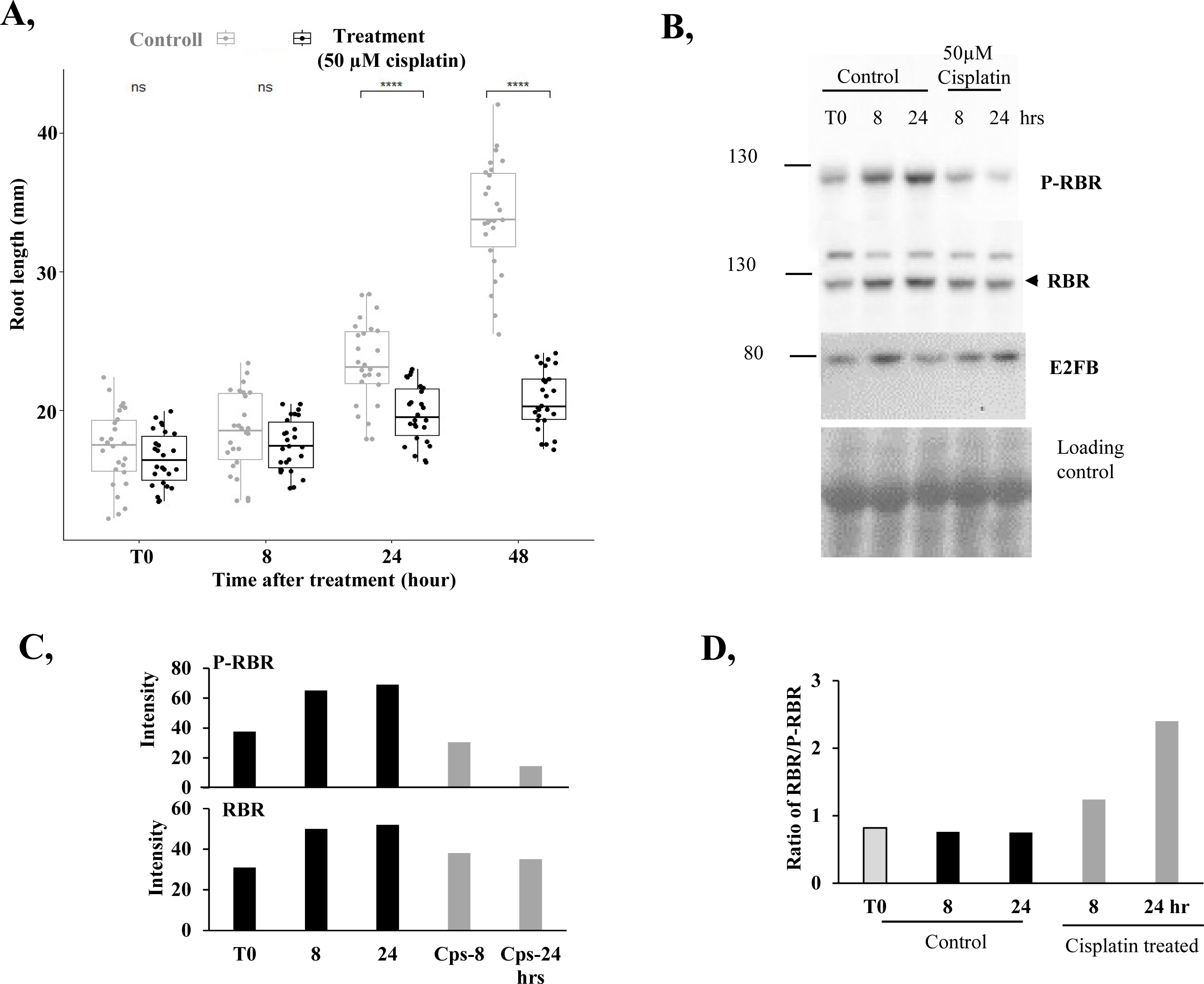
DNA damage inhibits root growth and decreases the phosphorylation level of RBR at the S911 site. (**A**) Root length of *A. thaliana* seedlings was measured in the presence of cisplatin (50 µM) and compared to the non-treated control roots. Data are average +/-standard deviation (n = 3 biological replicates; N= 10 samples in each), ns means non-significant changes, ****P ≤ 0.001 indicates statistically relevant differences between the mutant and the WT (two-tailed paired t test between the mutant and the WT). (**B**) Western blot analysis demonstrates P-RBR, RBR, and E2FB levels in both non-treated and cisplatin treated seedlings 4 days after germination (DAG; T0) and 8 and 24 hours after incubation with the drug. The intensity (**C**) and the ratio (**D**) of P-RBR and RBR signals was determined using the ImageJ software.

**Figure 3.**
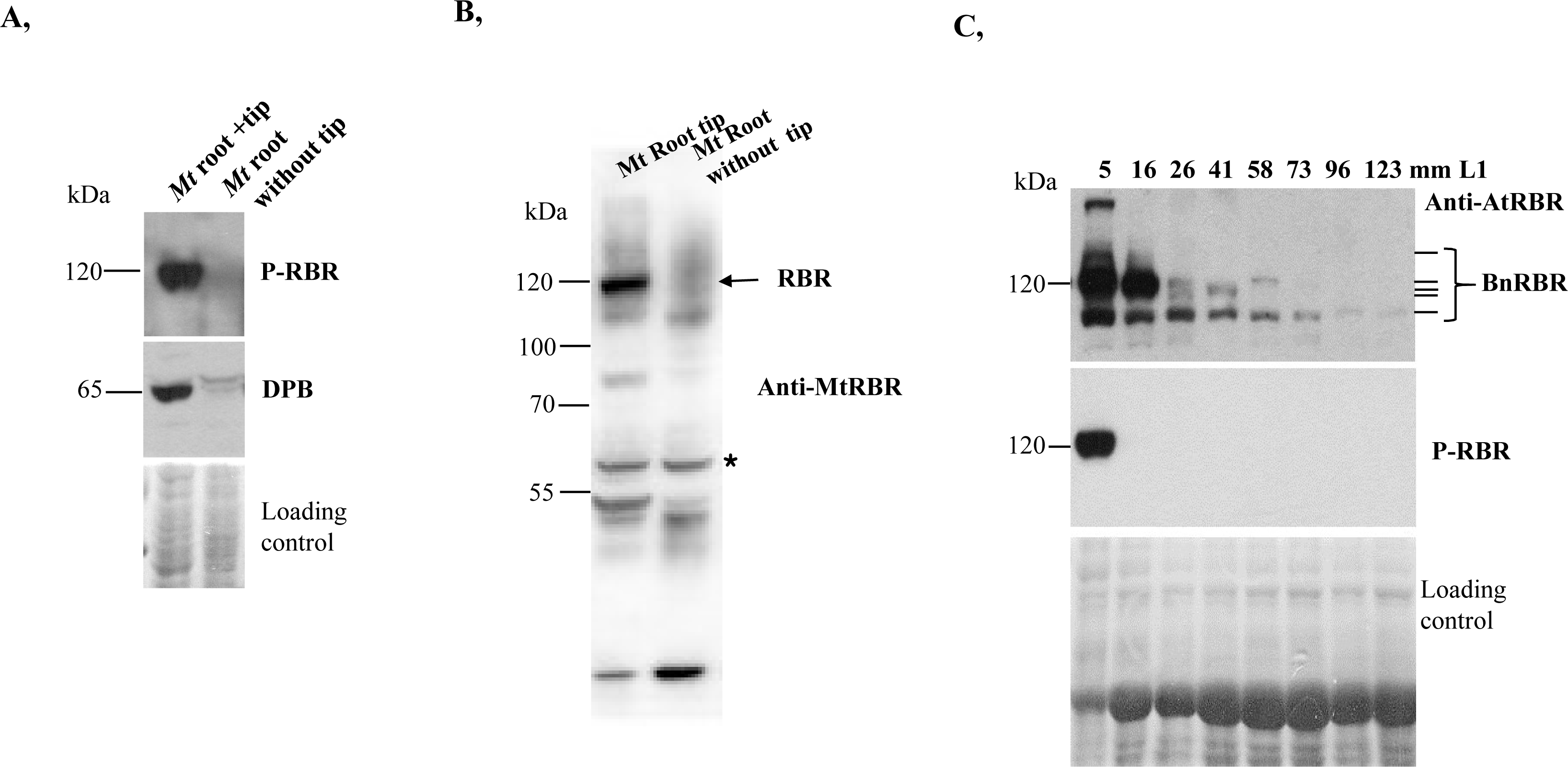
Phosphorylation of plant RBR at the conserved serine site corresponding to the animal Rb’s S807 increases in plant organs enriched with proliferating cells. (**A**) Western blot analysis demonstrates P-RBR^S807/811^, and DPB accumulation levels in either the root tip or elongation root section of *M. truncatula* root. Ponceau-stained membrane-bound proteins served as loading control. Molecular weight markers are indicated on the left side. (**B**) *M. truncatula* root proteins were also analysed via Western blot using an N-terminal specific *M. truncatula* MtRBR antibody. Arrow indicates specific protein. Asterisk denotes a protein indicating equal loading. (**C**) Western blot analysis reveals RBR and P-RBR levels in developing B. napus first real leaf harvested with various diameters as indicated in millimetre (mm). Ponceau-stained membrane proteins served as loading control. Molecular weight markers are indicated on the left side. The black lines, accompanied by a brace on the right side, denote proteins corresponding to the molecular size of potential various Brassica RBR proteins.

The phosphorylation site corresponding to Arabidopsis 911S is conserved and present in numerous plant RBR sequences (Wang et al., 2014), suggesting that its phosphorylation might function similarly in various plant species. To investigate whether the phosphorylation of RBR at this specific site also exhibits developmental regulation in other species, we analysed its pattern in the *Medicago truncatula* root and in developing *Brassica napus* leaves using both phospho-site and RBR-specific antibodies in western blot assays (Fig. 3). In *M. truncatula*, the root tip, comprising the root meristematic region, is enriched with dividing cells, as visualized by EdU-labelling marking root cells in S-phase (Suppl. Fig. 1A). Root tips and less proliferative root elongation and differentiation regions of *in vitro* grown 3-day-old *M. truncatula* seedlings were dissected and harvested separately. The phospho-site-specific signal was prominent in the proliferative root tip sample, yet it was undetectable in the less proliferative root section (Fig. 3A). The proliferation specific DPB protein demonstrated a similarly elevated expression level in the root tip, whereas its expression was markedly diminished in the roots devoid of the tip (Fig. 3A). Interestingly, the expression level of the *M. truncatula* MtRBR protein at its expected size was exclusively detectable in the root tip samples (Fig. 3B). To confirm that the P-RB^807/811S^ antibody indeed recognized the phosphorylated form of MtRBR at the corresponding 931S site, the protein was first immunoprecipitated using the phospho-specific animal RB antibody and then blotted with the AtRBR antibody (Suppl. Fig. 1B). The results confirmed the specificity of the antibody in the *M. truncatula* root. RBR- and phospho-site-specific signals were also examined in the developing first leaf pairs of *Brassica napus* using an immunoblot assay. *B. napus* first leaves were collected at various stages of development, ranging from very small to significantly larger sizes (between 0.5 and 12 cm in diameter) representing proliferative and elongating leaves, respectively. Proliferating cells in mitosis with condensed chromosomes were exclusively detected in the epidermis of the small-sized leaf under a DIC microscope (Suppl. Fig. 1C). The phospho-site-specific RBR antibody signal was detected only in the samples of the smallest leaves in agreement with their high proliferative activity (Fig. 3C). The *B. napus* RBR protein, like its counterpart in Arabidopsis, exhibited developmental regulation, being most abundant in the small-sized leaf and gradually decreasing in the elongating and differentiating leaf (Magyar et al., 2012; Őszi et al., 2020).

Collectively, these data support that phosphorylation of plant RBRs at sites corresponding to the 911S site of Arabidopsis RBR predominantly occurs in proliferating plant organs and meristems.

### P-911S in conjunction with other phosphorylated sites prevents RBR binding to E2Fs and other DREAM components

Peptides phosphorylated at the 911S site were not present in the Arabidopsis RBR proteins co-immunprecipitated by E2Fs (Table 1). The anti-P-RB^807/811S^ antibody has been demonstrated to efficiently immunoprecipitate P-RBR^911S^ from Arabidopsis (Magyar et al., 2012). Thus, this antibody was employed to immunprecipitate P-RBR^911S^ from one-week-old Arabidopsis seedlings expressing RBR-GFP, to examine whether the immunoprecipitated RBR is mono- or multi-phosphorylated. Mass spectrometry of the immunoprecipitate indicated that P-RBR^911S^-GFP could be phosphorylated at any of the 13 previously identified CDK sites. This suggests the prevalence of multi-site phosphorylation of the P-911S-containing RBR (Table 1). An additional IP was performed on identical samples using an anti-GFP matrix, followed by mass spectrometry analysis. All 13 CDK sites were again detected to be phosphorylated corroborating the previous results. This immunoprecipitate may encompass all forms of RBR, comprising a combination of various mono-, multi-, or non-phosphorylated variants. For the sake of clarity, the proteins precipitated using anti-GFP for RBR-GFP- or anti-AtRBR for endogenous RBR will be collectively referred to as RBR and the anti-P-RB^807/811S^-precipitated Arabidopsis RBR subpopulation will be designated as P-RBR^911S^ representing all phosphorylated RBR forms carrying P-911S.

The above immunoprecipitates were also investigated for interacting protein partners of RBRs. The E2FB, DPB, and the DREAM component ALWAYS EARLY 3 (ALY3, the plant homolog of the animal Lin 9) were readily identified among the proteins associated with RBR-GFP (Lang et al., 2021; Suppl. Data 2). In contrast, neither E2Fs nor DPs, nor the DREAM component ALY3, were observed in the anti-P-RB807/811S-precipitated RBR^911S^-GFP or RBR^911S^ samples (Suppl. Data 2). Corroborating the Arabidopsis P-RBR^911S^-related data, phosphorylated P-RBR^908/931S^ forms from Medicago root and Brassica leaves, respectively, were not observed in complex with E2F, DP, or the other DREAM component ALY3 (Suppl. Data 3). It is therefore plausible to hypothesize that the phosphorylation of sites corresponding to 911S in AtRBR marks the inability of plant RBRs to form complexes with these cell cycle regulators. Our data also suggest that these RBR proteins are predominantly multi-phosphorylated, as their sequencing identified various phosphorylated peptides in addition to P-911S or its equivalents (Suppl. Table 1).

### An activated RBR-kinase in the *e2fab* double mutant highly phosphorylates RBR at the 911S site but not in E2FC-RBR complex

The data presented above indicate that phosphorylation of RBR at 911S is coupled to cell proliferation concentrated in meristems and young developing organs such as leaves and prevents binding to E2Fs. This observation supports the hypothesis that phosphorylation at this site in Arabidopsis and its corresponding site in other plant species is driven by CDKs and may be crucial for cell cycle entry. Indeed, overexpression of CYCD3;1 has been shown to lead to RBR elevated phosphorylation at 911S and hyperplasic growth, suggesting that ectopic CYCD3;1 promotes cell proliferation through RBR phosphorylation at the 911S site (Dewitte et al., 2003; Magyar et al., 2012). In contrast, ectopic expression of the CDK inhibitor KIP-RELATED PROTEIN 2 (KRP2) led to decreased phosphorylation of RBR at the 911S resulting in suppressed cell proliferation (Magyar et al., 2012). It has also been previously reported that in the *e2fabc* triple mutant RBR was found to undergo significant phosphorylation at the 911S site (Gombos et al., 2023). In the leaves of the *e2fabc* mutant cell proliferation is notably hyperactivated due its inability to control cellular quiescence requiring E2F and RBR containing repressor complexes (Gombos et al., 2023). Contrary to the triple *e2fabc* mutant, the double *e2fab* mutant did not show hyperplasia and leaf size is unaffected in this double mutant in comparison to the wild type (Gombos et al., 2023). Consistent with this observation, the S-phase regulatory gene, *PROLIFERATION CELL NUCLEAR ANTIGENE 1* (*PCNA1*), was found to be up-regulated exclusively in the *e2fabc* triple mutant leaf compared to the WT control and the *e2fab* double mutant (Suppl. Fig. 2A). Furthermore, we did not observe a significant difference in the number of S-phase cells even within the root meristem of WT and *e2fab* double mutant when examined under a confocal laser microscope utilizing the EdU incorporation assay (Suppl. Fig. 2B-D). Nevertheless, RBR was found to undergo significant phosphorylation at the 911S site in the *e2fab* double mutant and in the *e2fabc* triple mutant in a similar manner (Fig. 4A). To elucidate the molecular mechanism underlying this surprising observation, we investigated the expression levels of *CYCD3;1*, *RBR*, and the F-Box protein *FBL17*, which downregulates KRPs at the protein level, during leaf development in the *e2f* mutants, comparing these levels to those in the WT (Fig. 4B-D). It is to be noted that *CYCD3;1*, *RBR*, and the *FBL17*, are all direct targets of the E2Fs (Zhao et al., 2012; Noir et al., 2015; Bouyer et al., 2018; Őszi et al., 2020; Gombos et al., 2023). It was found that *RBR* and *FBL17* were up-regulated in both mutants in correlation with the increased phosphorylation of 911S in RBR. This finding supports the hypothesis that the E2FA and E2FB regulated FBL17 protein, by facilitating the degradation of KRP proteins, activates RBR-kinases for phosphorylation at the 911S site (Noir et al., 2015; Gentric et al., 2020). *CYCD3;1* expression was significantly upregulated in young leaves, especially in the highly proliferative *e2fabc* mutant, exhibiting correlation with the rate of cell proliferation. However, its expression did not correlate with P-RBR^911S^ levels, suggesting that it may not be directly involved in the enhanced phosphorylation of RBR at 911S throughout the leaf development of *e2f* mutants.

**Figure 4.**
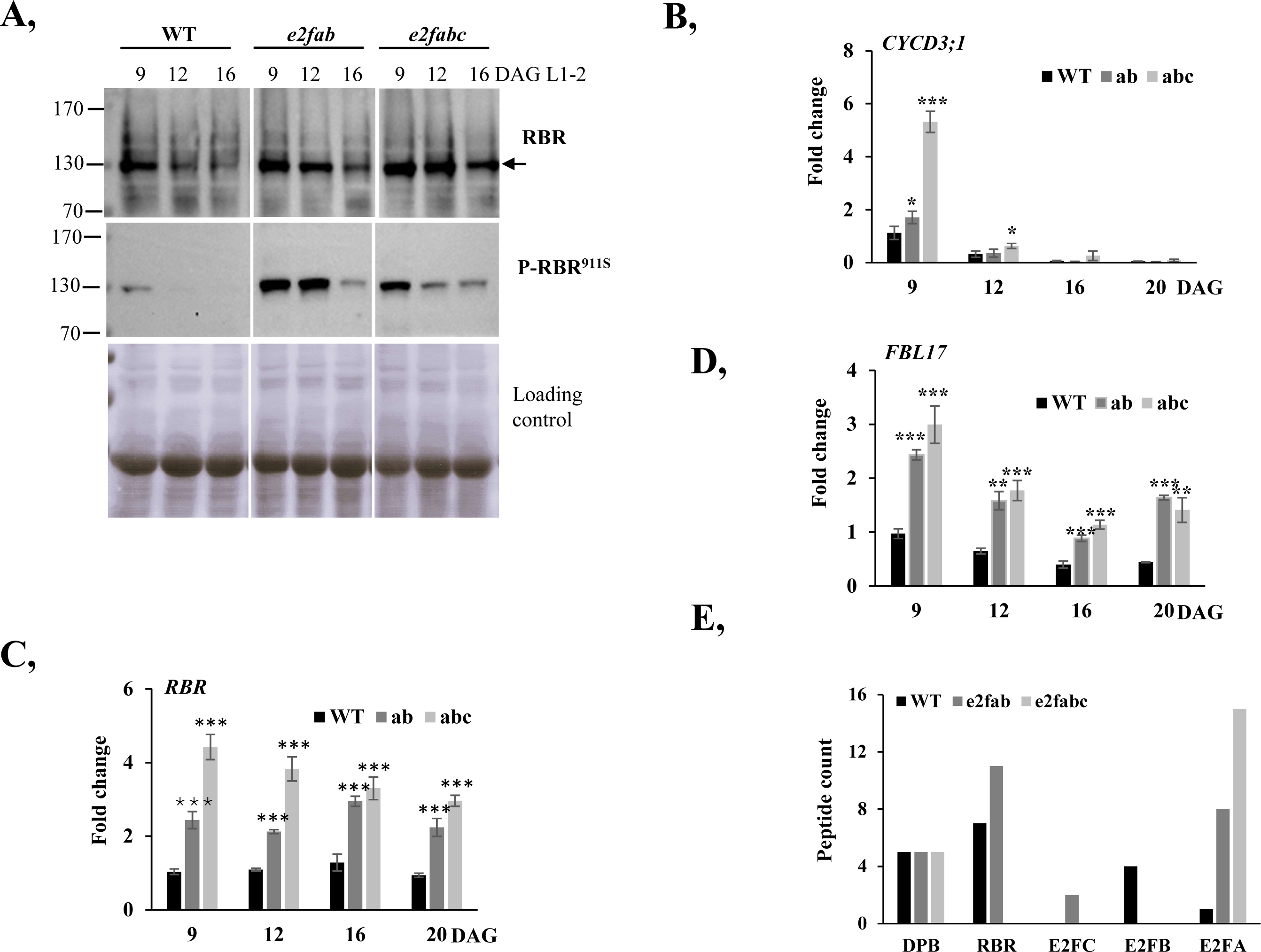
RBR phosphorylation at the serine 911 site in *e2f* mutants. (**A**) Western blot analysis demonstrates RBR presence and its phosphorylation level at 911S during leaf development of wild type, *e2fab* and *e2fabc* mutant seedlings 9, 12, 16 days after germination (DAG). Arrow marks RBR protein. Ponceau-S-stained membrane bound proteins demonstrate equal loading of protein samples. Molecular weight marker is indicated on the left side. (**B-D**) Expression of CYCD3;1 (B), RBR (**C**) and FBL17 (**D**) genes was monitored during leaf development of WT, *e2fab* and *e2fabc* lines at the indicated days after germination (DAG) utilizing qRT-PCR. Values represent fold changes normalized to the value of the relevant transcript of the wild type at 9DAG, which was set arbitrarily at 1. Data are means +/- sd., n=3 biological repeats. ***P<0.001 (two-tailed, paired t test between the WT and the mutant lines at a given time point). (**E**) Total peptide counts of DPB, RBR, E2FC, E2FB and E2FA proteins detected after mass spectrometry analysis of anti-DPB immunoprecipitated complexes in leaf samples of WT, *e2fab*, and *e2fabc* lines at 9DAG.

The data show that in the absence of E2FA and E2FB, RBR is highly phosphorylated at 911S due to the activation of an RBR-kinase. Despite the heightened phosphorylation, there is no corresponding increase in cell proliferation, implying that RBR is not fully suppressed in the *e2fab* double mutant. This is likely due to the presence of E2FC-RBR complexes, which are presumably unaffected by this RBR kinase and are capable of efficiently repressing cell proliferation (Gombos et al. 2023). Previous research has demonstrated that DPB effectively associates with RBR in young developing leaves of the *e2fab* mutant through heterodimer formation with E2FC, but this association is absent in the *e2fabc* triple mutant (Gombos et al., 2023). To examine the interaction between E2F-DPB heterodimers and phosphorylated RBR forms in detail, protein complexes were immunoprecipitated using an anti-DPB antibody from the first leaf pairs of WT, *e2fab*, and *e2fabc* lines at the young proliferative developmental stage. Subsequently, the associated E2F and RBR proteins, along with the phosphorylation status of DPB-bound RBR, were analysed through mass spectrometry. Notably, the E2FC protein, which exhibits only weak expression in the WT, was detected in complex with DPB exclusively in the *e2fab* double mutant but not in the WT unlike E2FB (Fig. 4E; Lang et al., 2023, Gombos et al., 2023). Conversely, E2FA associated with DPB in all three lines, indicating that mutant, C-terminal truncated E2FA in *e2fa-2* retains its capacity to dimerize with DPB (Leviczky et al., 2019; Gombos et al., 2023). RBR was detected in complex with DPB in the WT and *e2fab* double mutant genetic backgrounds but not in the triple *e2fabc* one (Fig. 4E). Its level was highest in the *e2fab* double mutant leaf (Fig. 4E; Gombos et al., 2023). Among the CDK sites of RBR two specific sites, namely 375S and 885S (refer to Table 1), were identified as phosphorylated when it was in complex with DPB, whereas the 911S site was not.

In conclusion, the RBR associated with E2FC-DPB heterodimers in the *e2fab* double mutant remains unphosphorylated at the 911S site, suggesting that the RBR kinase responsible for phosphorylating RBR at this position is unable to repress RBR within its E2F-bound complexes and thus cannot promote cell cycle entry.

### The 911S site is accessible for CDKs only when RBR is not in complex with E2Fs-DPB

Based on the assumption that E2FC-RBR complexes remain stable at high P-RBR^911S^ levels in the *e2fab* mutant, we supposed that this site has varying accessibility for the CDK depending on whether RBR is in complex with E2F-DP or not. To test this hypothesis, we applied molecular dynamics (MD) simulations on RBR in both configurations (Fig. 5 and Suppl. Fig. 3). Monomeric RBR shows significantly greater flexibility near the 911S site compared to its complexed form (Suppl. Fig. 3). Additionally, the 911S site is masked in the RBR-E2FB-DPB complex, whereas it remains exposed in the monomeric state (Fig. 5). We assessed flexibility in two additional CDK site regions known to be involved in E2F-binding (Zamora-Zaragosa 2024): around 406T and in the linker region between the A- and B pockets. Both regions display substantial flexibility in the E2F-free and E2FB-DPB complexed states of RBR (Suppl. Fig 3). The simulations indicate that the 911S site in the RBR-E2F-DP complex is stable, with limited CDK phosphorylation accessibility (Fig. 5) unlike other CDK sites like 406T and 712S, which remain accessible for phosphorylation even when RBR is complexed with E2Fs-DPB (Suppl. Fig 4). Therefore, distinct phosphorylation events at various RBR sites may regulate its E2F-binding and release activities, thereby refining its function as a transcriptional repressor.

**Figure 5.**
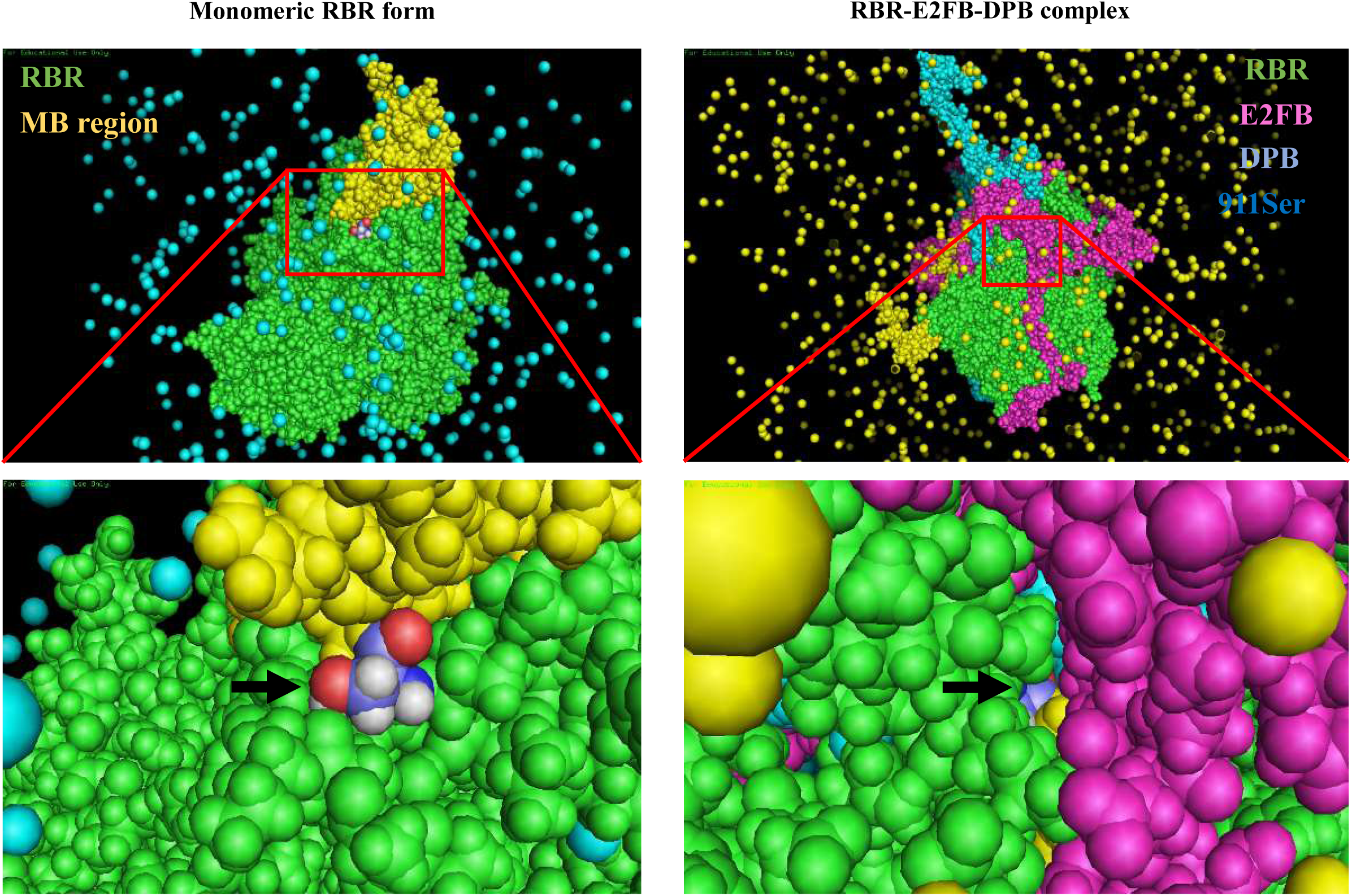
The predicted accessibility of the 911S site of free or E2FB/DPB-bound Arabidopsis RBR. Prediction of the three-dimensional protein structure of monomeric RBR and RBR-E2FB-DPB complex using Alphafold 3 (AF3) with the following colour code: green: RBR; pink: E2FB; light blue: DPB; dark blue-burgundy-white: side chain of 911 serine residue (911S). The arrow indicates the location of the 911S site within the monomeric RBR and when its complex with E2FB-DPB. Solvents, specifically water molecules, are depicted in blue on the left side and in yellow on the right side.

### Phosphorylation of RBR at the 911S site alters its function

We identified 77 high confidence interactors in the Arabidopsis P-RBR^911S^ pull down experiment (Suppl. Data 3). Interestingly, a significant number of interactors (28 out of the total 77) are associated with various RNA-binding functions. To further elucidate the different functional roles of proteins associated with RBR and P-RBR^911S^ forms, respectively, we compared the Gene Ontology (GO) enrichment analyses of their interactors using the ShinyGo 0.82 program (Ge et al., 2020; Fig. 6). Among the interactors of all RBR forms, the members of transcriptional repressor complexes like the DREAM (DRM) complex were prominently identified in the category of cellular components (Fig. 6A). Conversely, members of the transcriptional repressor complex were not present among the interactors of the P-RBR^911S^-containing subpopulation. Instead, these interactors were preferentially associated with the pre-ribosome large subunit precursor as cellular component (Fig. 6B). With respect to biological processes, all RBR forms were identified in association with proteins involved in protein transport or ribosome biogenesis. It appears, however, that members of the P-RBR^911S^ subpopulation are less involved in the transport function, suggesting that these interactions might necessitate the presence of E2Fs. Based on their interacting partners, the P-RBR^911S^ forms predominantly function in co-transcriptional and post-transcriptional regulations (Fig. 6B). The proteins interacting with the various P-RBR^908/931S^ forms of *Medicago truncatula* and *Brassica napus* were also found to be enriched in proteins exhibiting RNA-binding activity like that of Arabidopsis (Suppl. Data 3). Notably, in all three plant species, P-RBR forms were identified in complexes with proteins involved in ribosome biogenesis and protein translation (Suppl. Fig. 4).

**Figure 6.**
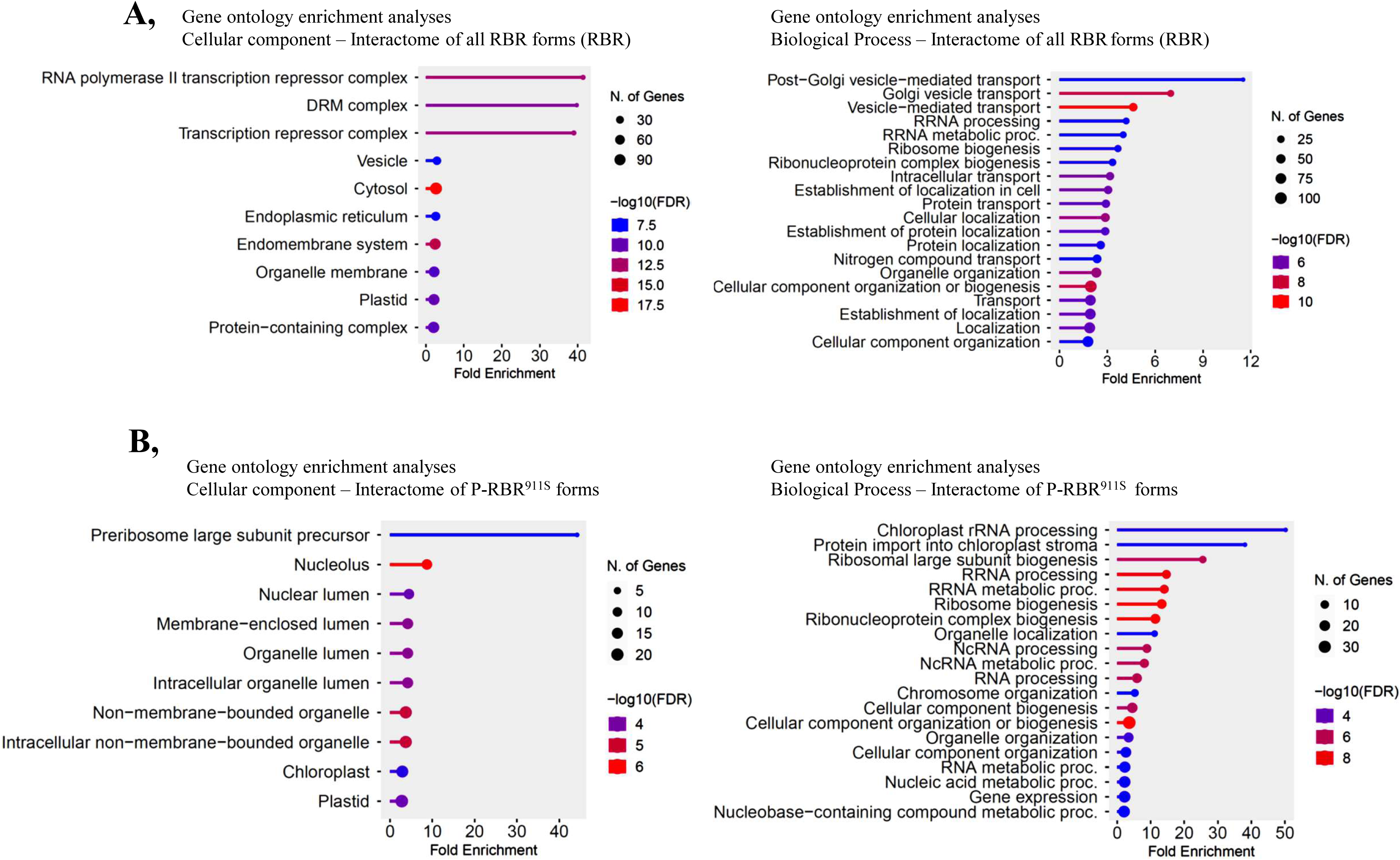
Gene ontology analysis of 911S phosphorylation-dependent interactors of Arabidopsis RBR. The gene ontology analysis of protein components associated with all RBR forms (**A**) and those where 911S is phosphorylated (**B**). The analysis was performed utilizing the ShinyGo 0.82 program. This analysis identifies proteins linked with RBR and P-RBR^911S^ that participate in various cellular components and biological processes, employing a false discovery rate (FDR) threshold of 0.05.

In conclusion, the (multi)phosphorylated P-RBR^911/908/931S^ forms of Arabidopsis, Medicago or Brassica could not be detected in complex with E2Fs, DPs, and DREAM components; however, they interacted with various proteins involved in ribosome biogenesis and mRNA splicing. This suggests that the phosphorylation of the site corresponding to 911S in Arabidopsis in plant RBRs (likely multi-phosphorylated), modifies the protein’s biological function from transcriptional regulation to post-transcriptional/translational control.

## Discussion

The research presented herein experimentally verified 13 phosphorylation sites among the 16 *in silico* predicted conserved CDK recognition motifs in various plant RBR proteins (Fig. 1). This observation aligns with the regulation of plant and animal RBR/RB proteins at similar phosphorylation sites suggesting a conserved cell cycle control mechanism across kingdoms. This control centres on RB’s ability to interact with and repress the activity of E2F/DP cell cycle-controlling transcription factor dimers, which is controlled by CDK-mediated phosphorylation of RB.

Interestingly, phosphorylation at multiple CDK sites was found not to inhibit the plant RBR’s interaction with E2Fs (Table 1). Nine distinct phosphorylated RBR forms retained the ability to bind E2Fs and DPB. Among these sites, the phosphorylation of five specific ones (385S, 389S, 406T, 685S, and 712S) is implicated in modulating the interaction between the RBR’s pocket region and the transactivation domain of E2F (Konagaya et al., 2024). The data imply that RBR phosphorylation at these sites is unable to fully disrupt the complex. A possible explanation is that RBR with its C-terminus can remain in complex with E2Fs through their MARKED box (MB) region. While the MB region is highly conserved in the plant E2F-DPs, its role in association with RBR binding has not yet been confirmed.

In animal cells, the RB protein was shown to be unphosphorylated in the G0 phase, monophosphorylated at any of the 14 CDK sites in the G1 phase, and hyperphosphorylated at all sites before the G1/S cell cycle phase transition (Narashima et al., 2014; Sanidas et al., 2019). While hyperphosphorylation of RB at the restriction point is required for the release of the E2F transcription factors from repression by RB, the 14 monophosphorylated RB isoforms were found to retain differential binding affinity towards the various E2F proteins in the G1 phase. Furthermore, these monophosphorylated isoforms of RB have been shown to associate with distinct protein partners to control various cellular processes (Sanidas et al., 2019). Recent research has indicated that the phosphorylation of animal RB occurs in two phases. The first phase involves the phosphorylation of sites that disrupt the interaction between the pocket domain of RB and the transactivation region of E2Fs (Konagaya et al., 2024). However, complete dissociation of RB from E2F requires further phosphorylation at the C-terminus, including the serine 807/811S sites (the site corresponding to 911S in Arabidopsis RBR). In our investigation, we could not discriminate the mono- or multi-phosphorylated RBR forms. It is possible that, similarly to animals, the P-RBR forms associated with E2Fs are monophosphorylated at various sites. However, the RBRs immunoprecipitated using an antibody specific for the phosphorylated RBR at 911S site were found to be phosphorylated in combination with other sites and were not forming complex with E2Fs and DPs, suggesting that multi-site phosphorylation including P-RBR^911S^ suppresses RBR’s E2F repressor function. Multi-phosphorylation has similar effect on the animal RB (Rubin SM; 2013; Knudsen and Wang 1996). In Arabidopsis, the phospho-mimicking mutation of the single 406T site exhibited a notable influence on meristem size depending on the phosphorylation status of other CDK sites, supporting the significance of combinatorial RBR phosphorylation in plants (Zamora-Zaragoza et al., 2024).

Structural modelling shows that the RBR’s 911S site is buried in the RBR-E2F-DP complex and becomes accessible for phosphorylation by CDKs only when the RBR is unbound to E2F-DPs (Fig. 5). In contrast, the 406T site of the RBR protein, implicated in the control of E2F-release, remains accessible to CDKs even when the protein is in complex with E2F-DPs (Fig. 5 and Suppl. Fig. 3). This observation suggests that phosphorylation of 911S (or its equivalent in other plant species) may influence the E2F binding activity of free RBR, but it does not affect the action of the already established transcriptional repressor RBR-E2F complexes. Studying P-RBR^911S^ levels in association with contrasting cell proliferation in *e2fab* and *e2fabc* mutants also support this view (Fig. 4). The elevated abundance of P-RBR^911S^ cannot destabilize the existing E2FC-DPB-RBR complexes in the *e2fab* mutant, allowing the establishment of proper cellular quiescence. Otherwise, the *e2fab* mutant should exhibit hyperplastic growth similarly to the *e2fabc* mutant (Gombos et al., 2023). It also agrees with the rapid decline in RBR phosphorylation at 911S in response to cisplatin treatment (Fig. 2). RBR is known to repress E2F activity in stress responses and DNA damage repair pathways (Horvath et al., 2017; Biedermann et al., 2017; Lang et al., 2021; Nisa et al., 2023). In agreement, the formation of DREAM complex in seedlings was observed at higher level under conditions of DNA damage (Lang et al., 2021). To allow RBR-E2F interaction under these conditions, the 911S site must also be unphosphorylated. The rapid accumulation of non-phosphorylated RBR at the 911S in response to DNA damage suggests that RBR may act as a phosphorylation state-dependent molecular switch, quickly halting cell cycle progression to allow for DNA repair. Therefore, we propose that plant RBR in complex with E2F-DP heterodimers may undergo sequential phosphorylation, analogous to the process observed in animal cells, where the RB at the 807/811S sites are phosphorylated during the second phase rather than the initial phase of E2F-release (Kim et al., 2022; Chung et al., 2019; Konagaya et al., 2024). One could suppose that phosphorylation at the 911S site in Arabidopsis (the site corresponding to 807/811S in human Rb) could be part of RBR hyperphosphorylation required for full RBR-E2F complex dissociation and the prevention of its reassociation. Therefore, the P-911S site must be dephosphorylated in response to DNA damage as well as during the G1 phase to allow RBR binding to E2F/DPs. Research has shown that RBR from *Medicago sativa* undergoes phosphorylation at the site corresponding to 911S when cells enter the cell cycle during the S-phase. This phosphorylation remains consistently high throughout the entire cell cycle but decreases during the G1 phase when RBR is in complex with E2Fs (Ábrahám et al., 2011)

The interaction of P-RBR^911S^ forms is not only inhibited with E2Fs and other DREAM components, but also with transport proteins linked to the Golgi apparatus and vesicles. This is evidenced by the absence of proteins with these functions from the interactome of P-RBR^911S^ forms, while they are enriched in the interactome of all RBR forms (Fig. 6). In contrast, the (multi)phosphorylated P-RBR^911S^ specifically interacts in Arabidopsis with proteins possessing RNA-binding activity and involved in biological processes such as ribosome biogenesis, RNA processing, and ribonucleoprotein complex assembly (Fig. 6). The RBR proteins of *Medicago truncatula* and *Brassica napus* that were phosphorylated at the corresponding sites were also found to be preferentially associated with proteins involved in ribosome assembly, and translation (Suppl. Fig. 4). These findings imply that the phosphorylation of RBR at 911S, likely in conjunction with other sites, could modify its role from transcriptional to post-transcriptional regulation. Earlier studies have already indicated the involvement of RBR in translational control (Lokdarshi et al., 2020). The TARGET OF RAPAMYCIN (TOR) kinase pathway, a key regulator of growth, modulates translation and ribosome biogenesis (González and Hall, 2017; Ahmad et al., 2019). In plants, TOR signalling has been shown to regulate E2Fs and RBR, while also enhancing the activity of EBP1, a driver of plant growth (Henriques et al., 2010; Xiong et al., 2013; Wu et al., 2019; Van Leene et al., 2019; Horváth et al., 2006). EBP1 operates within complexes containing RNA-binding proteins that are integral to the regulation of protein synthesis. It has been shown to mitigate the cell cycle inhibitory function of RBR, promoting growth (Horváth et al., 2006). Therefore, RBR and EBP1 are antagonistic. They exert their antagonism by interacting with a shared set of partners implicated in ribonucleoprotein complex formation (Lockdarshi et al., 2020). Further research is necessary to elucidate the mechanisms by which the P-RBR^911S^ forms, in conjunction with RNA-binding proteins, affect protein translation and splicing. Additionally, it is crucial to understand how these RBR-associated post-transcriptional regulatory processes contribute to cell cycle control and meristematic activity. Furthermore, RBR and E2FB are in the same protein complex not only in mitotic but also in post-mitotic cells (Őszi et al., 2020; Lang et al., 2021; Hamid et al., 2025). In post-mitotic cells, these complexes might regulate functions that are unrelated to the cell cycle but still can be under the control of RBR’s phosphorylation state.

There is limited understanding of the temporal regulation of RBR phosphorylation during the plant cell cycle. Our results demonstrate that phosphorylated RBR forms can complex with E2Fs even during the G1 phase. Only inhibitory multi-phosphorylation facilitates the release of E2Fs from RBR as cells progress into the cell cycle. This multi-phosphorylation, however, does not initially include the phosphorylation of the 911S (or equivalent) site (Fig. 7A). The subsequent phosphorylation of E2F-free RBR at the 911S site likely has a role in inhibiting the re-association of RBR with E2Fs while promoting the RBR’s interaction with ribosomal and splicing proteins, thereby contributing to the post-transcriptional regulation of the cell cycle (Fig. 7B). Whether this RBR-mediated regulation positively or negatively influences the processes of ribosome assembly and protein translation needs further investigation. The specific CDK complex responsible for phosphorylating RBR at the 911S site remains unidentified; however, the data suggest that these complexes may be the CDKs inhibited by the KRPs. Nevertheless, we propose that distinct cyclin-dependent kinases (CDKs) target RBR depending on whether it is in complex with E2Fs or not; the former releases E2Fs from RBR-binding, while the later inhibits the free RBR’s ability for (re)making complex with E2Fs and DREAM components (Fig. 7C). We suggest that the initiation of cell cycle entry necessitates the release of E2Fs from the repression exerted by RBR. Simultaneously, the unbound RBR must undergo multi-phosphorylation to prevent its ability to bind E2F.

**Figure 7.**
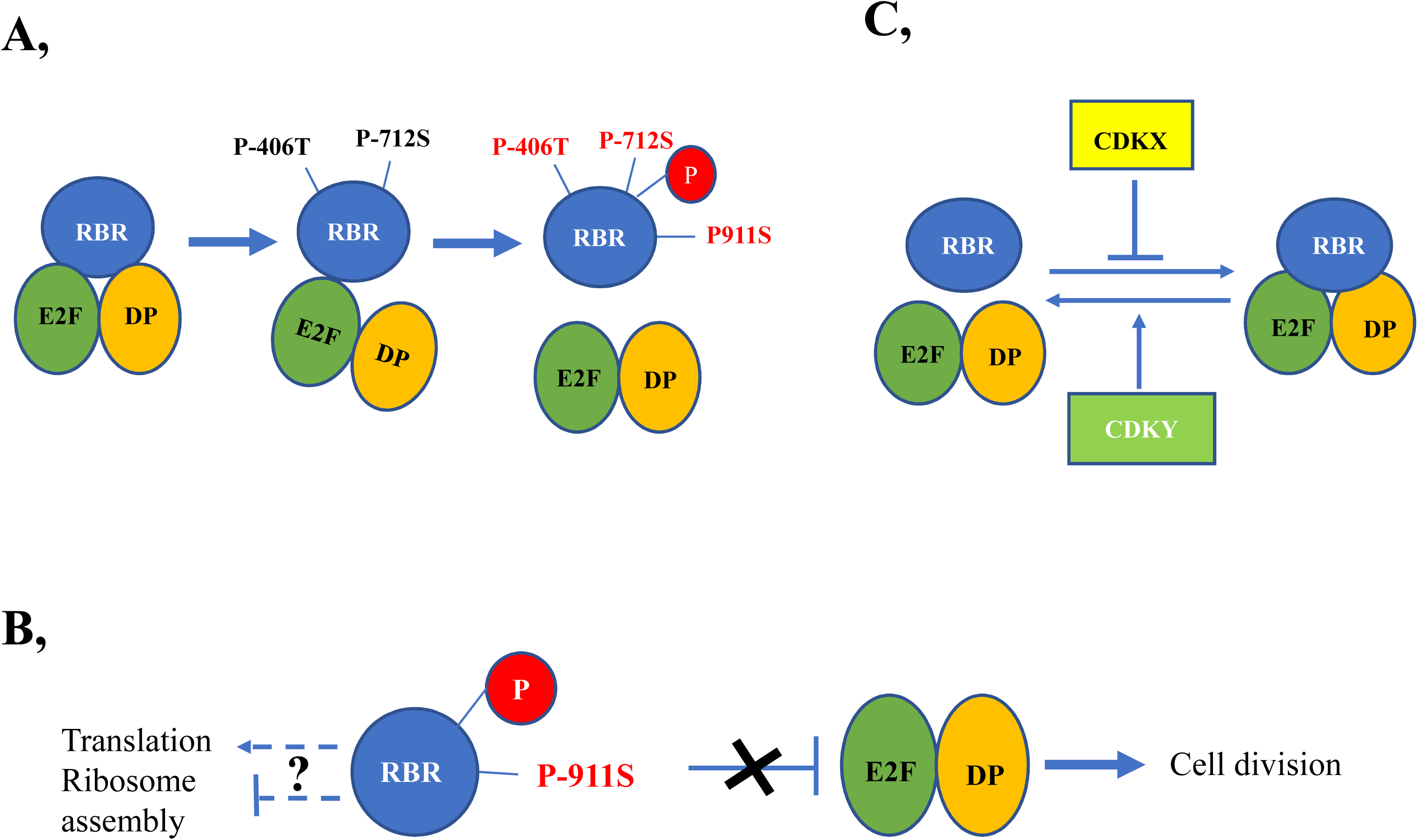
A model to explain the role of 911S phosphorylation in RBR function. The function of the RBR transcriptional repressor is modulated by at least two phosphorylation-based mechanisms. (**A**) Initially, the RBR-E2F-DP complex is suggested to undergo phosphorylation at sites such as 406T and 712S, which are accessible to CDKs, thereby weakening the interaction between RBR and the E2F-DP. However, subsequent phosphorylation at the C-terminal, including the 911S site, results in the irreversible disruption of the complex till the next G1 phase. RBR is likely to be multiphosphorylated at several sites beside those indicated by amino acid numbers. This is indicated by the capital P in red circle. (**B**) In the multiphosphorylated state, involving P-911S, the binding capacity of RBR to E2F-DP is inhibited. These RBR forms therefore are unable to contribute to the E2F/DP-mediated control of cell proliferation or quiescence. However, the binding of RBR to proteins involved in post-transcriptional processes such as translation and ribosomal assembly is facilitated by the multiphosphoprylation of RBR involving 911S. RBR may positively or negatively influence these post-transcriptional processes as indicated by a question mark. (**C**) It is likely that specific CDKs (marked as CDKX and CDKY) regulate these processes at different sites within the RBR protein.

The current findings advance our comprehension of RBR regulation in plants. They elucidate a more complex understanding of the influence of phosphorylation on RBR function, suggesting that distinct phosphorylation patterns may refine not only the transcriptional inhibitory function of RBR but also its potential regulatory role in the modulation of translation and transport. In conclusion, this study provides additional data on RBR phosphorylation in plants, revealing a sophisticated regulatory system that extends beyond a simple on/off control of RBR activity.

## Materials and Methods

### Plant material and growth conditions

*Arabidopsis thaliana* ecotype Columbia wild-type and transgenic seeds were sterilized using commercial bleach, subsequently re-suspended in sterile water, and subjected to cold treatment at 4 °C, in darkness for 2 days (Clough and Bent, 1998). Unless otherwise specified, the plants were cultivated under continuous light at 22 °C in vitro on half-strength germination medium with a light intensity of 100 μmol mE^−2^ s^−1^ light intensity. The first leaf pairs of the wild-type or the T-DNA insertion double *e2fab* (Heyman et al., 2011) and the triple *e2fabc* (Gombos et al., 2023) mutants were cultivated *in vitro* were harvested 9–20 days after germination (DAG), flash frozen, and stored at −80 °C. *Medicago truncatula* seedlings were grown under a 16-h light/8-h dark photoperiod at 22 °C in vitro on germination medium. The initial true leaf of *Brassica napus* seedlings, cultivated under a 16/8 hour light-dark cycle, was harvested at various sizes ranging from 5 mm to 123 mm, rapidly frozen under liquid nitrogen, and stored at −80°C.

### EdU staining of Medicago root explants and Arabidopsis roots

For the EdU incorporation assay, three-day-old *Medicago truncatula* wild-type root explants, including the root tips, were cultivated in half-strength liquid MS medium containing 10 μM EdU (Click-iT Alexa Fluor 647 Imaging Kit; Invitrogene) and incubated for three hours. In the case of Arabidopsis, five days-old WT and *e2fab* mutant were placed into half-strength liquid MS containing 5 μM EdU and incubated for 30 min. Subsequently, the root samples and the seedlings were treated as previously described (Gombos et al., 2023), and the roots were stained with Hoechst solution. Observations were conducted using a confocal laser microscope.

### Cisplatin treatment and root growth assay

Arabidopsis seeds were germinated on vertical plates containing ½ MS medium and cultivated under continuous light conditions at 22°C for five-day post-germination. Subsequently, the seedlings were then transferred to plates, either supplemented with 50 µM cisplatin (Merck, Germany) or without it. The plates were scanned alongside a ruler, and root length was digitally measured at the initial time point (T0) and 8, 24 and 48 hours thereafter using ImageJ software.

### Immunoblotting, and immunoprecipitation and generating antibody against *Medicago truncatula* MtRBR

Immunoblotting and immunoprecipitation assays were carried out as described earlier (Gombos et al., 2023). Primary antibodies used in these assays were anti-phospho-specific RB (Ser^807/811^) rabbit polyclonal antibody (Cell Signalling Tech), chicken anti-Arabidopsis C-terminal specific RBR antibody (Agrisera), mouse anti-Arabidopsis N-terminal specific RBR antibody (Hamid et al., 2025), rabbit anti-DPB antibody (Umbrasaite et al., 2010), rabbit anti-E2FB (Magyar et al., 2005).

For the *Medicago truncatula* RBR antibody production (MtRBR), the N-terminal domain of RBR (encoding the first 381 amino acids) was cloned into the expression vector pET28a to BamHI-SalI sites which provides an N-terminal 6X histidine tag.

For cloning the MtRBR N-terminal coding region we used cDNA synthetized with M-MLV Reverse Transcriptase (Thermo Scientific #28025013) and oligo(dT)_25_ as priming from 5 µg total RNA of 10 DAG old *M. truncatula A17* root tips grown under normal conditions. Than RBR N-terminal region (1143 nd) was amplified with Takara PrimeSTAR Max Polimerase with the following primers: MT-RBR-alpha-5’-BamHI: 5’-cgcggatccATGAGTCCTTCAGCTGAAACGG-3’ and MT-RBR-alpha-3’-SalI: 5’-acgcgtcgacCATCATCAAATCATATTTCCTC-3’ according to the manufacturer’s manual (https://www.takarabio.com/documents/User%20Manual/R045A/R045A_e.v1510Da.pdf).

Protein expression construct was transformed into *Escherichia coli* ArcticExpress cells. Induction and purification were done under native conditions as described in the AGILENT handbook (https://www.agilent.com/cs/library/usermanuals/Public/230191.pdf).

In short, a preculture was grown overnight in liquid Luria-Bertani (LB) medium at 37°C in 200 mg/L gentamycin and 50 mg/L kanamycin. The next day, the culture was transferred and diluted 1:100 in LB without antibiotics at 28 °C until an OD of 0.4 was reached. Then the culture was shifted to 10°C, adapted for 1 h, and induced with IPTG (1 mM final concentration). Than the cells were incubated for 24H at 10 °C with shaking 200-250 rpm. The cells were harvested (centrifugation for 15 min at 10,000 rpm) when they reached an OD of 1.5. For the subsequent steps, the cells were kept at −80 °C. Protein expression was analysed from 20uL of induced cultures by SDS-PAGE prior to protein purification.

For protein purification the pellet was re-suspended first in 2.5 mL resuspension buffer (50 mM Na-phosphate, 300 mM NaCl, and 10 mM imidazole) per 100 mL culture and subsequently centrifuged at 12,000 rpm for 5 min. Than the supernatant was discarded and the pellet was re-suspended in 2.5 mL lysis buffer (50 mM Na-phosphate, 300 mM NaCl, 10 mM imidazole, and 4 μg lysozyme) per 100 mL of cell culture. After an incubation time of 1 h, the extract was centrifuged at 20,000 rpm for 30 min. His-tagged recombinant proteins from the supernatant were purified under native conditions by using gravity-flow columns with Thermo Scientific HisPur Ni-NTA Superflow Resin (#25214) according to the manufacturer’s user guide (https://assets.thermofisher.com/TFS-Assets/LSG/manuals/MAN0011700_HisPur_NiNTA_Resin_UG.pdf).

For polyclonal antibody production mice (6-8 week old Balb/c) were immunized biweekly (three times) with 100 µg of the native protein extract/immunization supplemented with Freud’s Adjuvant in 1:1 ratio [Complet Freud’s Adjuvant (Sigma, Cat. no.: F5881) for the first immunization and Incomplete Freud’s Adjuvant (Sigma, Cat. no.: F5506) for the 2^nd^ and 3^rd^ immunization]. One week after the 3^rd^ immunization total serum was harvested and supplemented with sodium-azide (0.01-0.04%) than stored at 4 °C.

### Quantitative RT-PCR

RNA was extracted from the plant material using the CTAB-LiCl method as described by Jaakola et al. (2001). RNA samples were treated with DNase1 (ThermoScientific #EN0521) according to the manufacturer’s protocol. cDNA was performed using 1 μg of RNA with ThermoScientific Reverse Transcription Kit (#K1691) using random hexamers, following manufacturer’s guidelines. A mock reaction without RevertAid enzyme was conducted to verify absence of genomic DNA contamination. RT-qPCR was performed with SYBR Green (TaKaRa TB Green Primer Ex TaqII, #RR820Q) as per manufacturer’s instructions, using a BioRAD CFX 384 Thermal Cycler (BioRAD) with parameters: 50 °C 2 min, 95 °C 10 min, 95 °C 15 s, 60 °C 1 min, 40 cycles, followed by melting point analysis. Each reaction was conducted in triplicate, and specificity was confirmed by a single peak in the melting curve. Data were normalised to average Ct value of two housekeeping genes (ACTIN2 and UBIQUITIN CONJUGATING ENZYME 18). Specified Primer sequences are detailed in Supplementary Table 2.

### Plant material and protein extraction for LC-MS/MS

Wild type *Arabidopsis thaliana* seedlings, RBR-GFP transgenic Arabidopsis lines, *Medicago truncatula* roots, and *Brassica napus* leaves were harvested, in addition to protein extracts prepared from E2F immunoprecipitations as described in Lang et al. (year). Approximately 150–200 seedlings per sample were flash-frozen in liquid nitrogen for 2 minutes. Tissue was disrupted in a TissueLyser (Qiagen; 50 Hz, 4 × 30 sec) with 6 mm stainless steel beads and sterile sand. Total proteins were extracted in 800 μL RIPA lysis buffer (150 mM NaCl, 1% IGEPAL CA-630, 0.5% sodium deoxycholate, 0.1% SDS, 50 mM Tris-HCl pH 8.0) supplemented with 1 mM DTT, 1 mM PMSF, protease inhibitor cocktail (Sigma), 3 mM pNPP and additional phosphatase inhibitors, as well as 1 μM MG132. Lysates were cleared by centrifugation (16,000 g, 15 min, 4 °C).

### Immunoprecipitation for LC-MS/MS

For phospho-RBR enrichment (P-RBR-IP), clarified extracts were incubated with Protein A-Sepharose beads coupled to anti-RBR antibody, or with anti-GFP magnetic beads (Miltenyi, MACS® Technology) for RBR-GFP transgenic material. Parallel experiments included DPB immunoprecipitations from Arabidopsis leaves (WT, *e2fab* and *e2fabc* mutants). Following incubation, beads were washed twice with RIPA buffer, five times with PBS, and once with ammonium bicarbonate (ABC) buffer. Bound proteins were reduced with 10 mM DTT, alkylated with 22 mM iodoacetamide, and on-bead digested with sequencing-grade trypsin (Promega) for 2 h at 47 °C. Digestion reactions were acidified, and one-sixth to one-eighth of each digest was loaded onto EvoTips (Evosep) for LC-MS analysis.

### Mass spectrometry analysis

Peptides were analyzed by nanoLC-MS/MS using an Evosep One (15 SPD) system online coupled to an Orbitrap Fusion Lumos mass spectrometer (Thermo Fisher Scientific) operating in positive ion mode. Survey scans were acquired in the Orbitrap (R = 120,000 at m/z 200) and multiply charged precursor ions were selected in data-dependent mode for HCD fragmentation in the linear ion trap. MS/MS spectra were acquired in the ion trap, while MS1 spectra were recorded in the Orbitrap.

### Data processing and phosphorylation site identification

Raw data were processed with Proteome Discoverer (v1.4) and searched against the SwissProt database (downloaded 2019/06/12, 560,292 entries) using Protein Prospector (v5.15.1). Search parameters: enzyme = trypsin (max. 1 missed cleavage), precursor mass tolerance = 5 ppm, fragment mass tolerance = 0.6 Da, fixed modification = carbamidomethylation (Cys), variable modifications = N-terminal acetylation, methionine oxidation, N-terminal pyroglutamate formation, and phosphorylation of serine, threonine, and tyrosine residues (max. 2 variable modifications/peptide). Protein and peptide identifications were accepted with minimum scores of 22 and 15, and maximum E values of 0.01 and 0.05, respectively. To further refine identifications, an additional database search was performed against the UniProt.random.concat database (downloaded 2022/07/20) with *Arabidopsis thaliana* species restriction (136,466 proteins), including high-confidence hits (protein score >50) from the SwissProt search. Phosphorylation site localization was validated based on site-determining fragment ions.

### Alfafolding model predictions and molecular dynamics simulations of different RBR forms

The models of the RBR and RBR-E2F-DP complexes were generated using the Alphafold 3 prediction via its website (Abramson, 2024). The structure of RBR monomer was based on the experimentally determined model of Homo sapiens Rb1 (RCSB PDB entry: 4ELJ; Burke et al., 2012), and the structure of the Arabidopsis complex was also based on the experimentally determined human E2F-DP-RBC (RCSB PDB: 2AZE; Rubin et al., 2005).

Input data for the molecular dynamics were generated using CHARMM-GUI (Jo, 2008), in a TIP3 water solvent, in the presence of potassium and chloride ions at a pH of 7. Molecular dynamics with GROMACS (Abraham, 2015) involved minimization and a 0.125 ns (125000 steps) equilibration. Three parallel simulations were run on each model variants up to 100 ns, with 2 fs time steps.

RMSF (root mean square fluctuations) values of the protein’s alpha C-atoms were calculated as indicators of the flexibility of certain segments of the polypeptide chain.

Amount of solvent atoms within a set radius of certain residues was calculated as the measure of the accessibility of the local molecule segment.

## Acknowledgements

We thank Gábor Steinbach (HUN-REN Biological Research Centre, Szeged, Hungary) for his assistance with microscopy. Additionally, we extend our appreciation to János Györgyey for his critical review of the manuscript. This research was supported for A.P-Sz by the EU Horizon 2020 grant No. 739593; KIM NKFIA 2022-2.1.1-NL-2022-00005. Additionally, Z.M and F.V-N received funding from the Hungarian National Research Funding (NKFI-139202), and together with P.K a grant from the HUN-REN Biological Research Centre, Szeged as a highlighted research theme (454004). M.G was supported by the Hungarian National Research Funding (NKFI-146566), and for A.F received funding from the Hungarian National Research Funding (NKFI-146386).

## Author contributions

Z.M., A.P-Sz., and F.V-N conceived the project and designed the experiments. A.P-Sz and F.V-N carried out most of the experiments. F.N., M.G., E.M., RSBH., P.K and Z.M, collected materials and carried out some experiments. Z.M., A. P-Sz., F.V-N and A.F wrote and revised the manuscript with input from all co-authors. All authors have read and approved the final manuscript.

