## Supplementary material for "Complex regulation of RETINOBLASTOMA-RELATED’s interactions with E2Fs via phosphorylation": Suppl. Figures and Table

### Slide 1
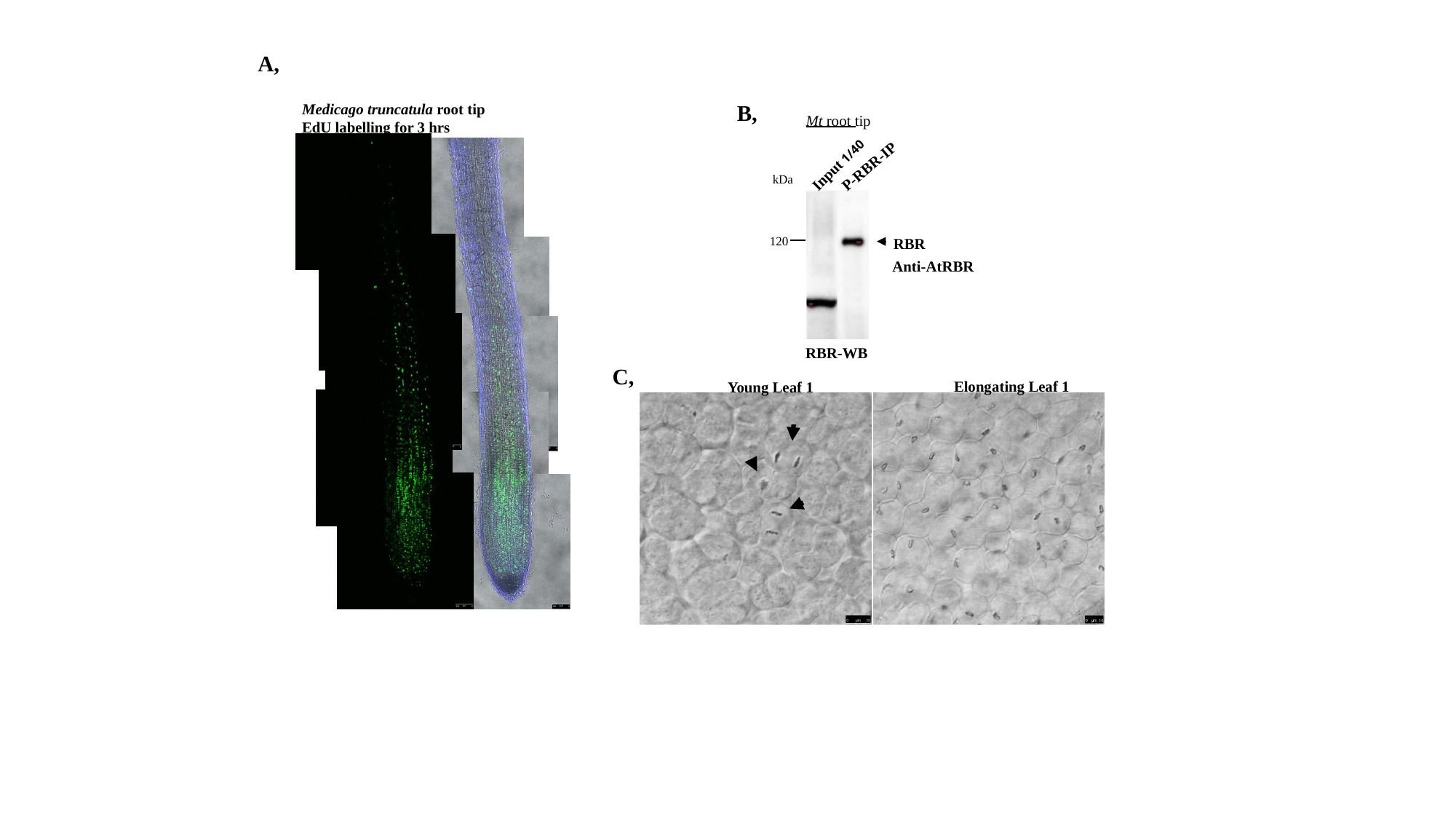

A,
B,
Mt root tip
Input 1/40
P-RBR-IP
120
Anti-AtRBR
RBR-WB
kDa
RBR
Medicago truncatula root tip
EdU labelling for 3 hrs
C,
Elongating Leaf 1
Young Leaf 1

### Slide 2
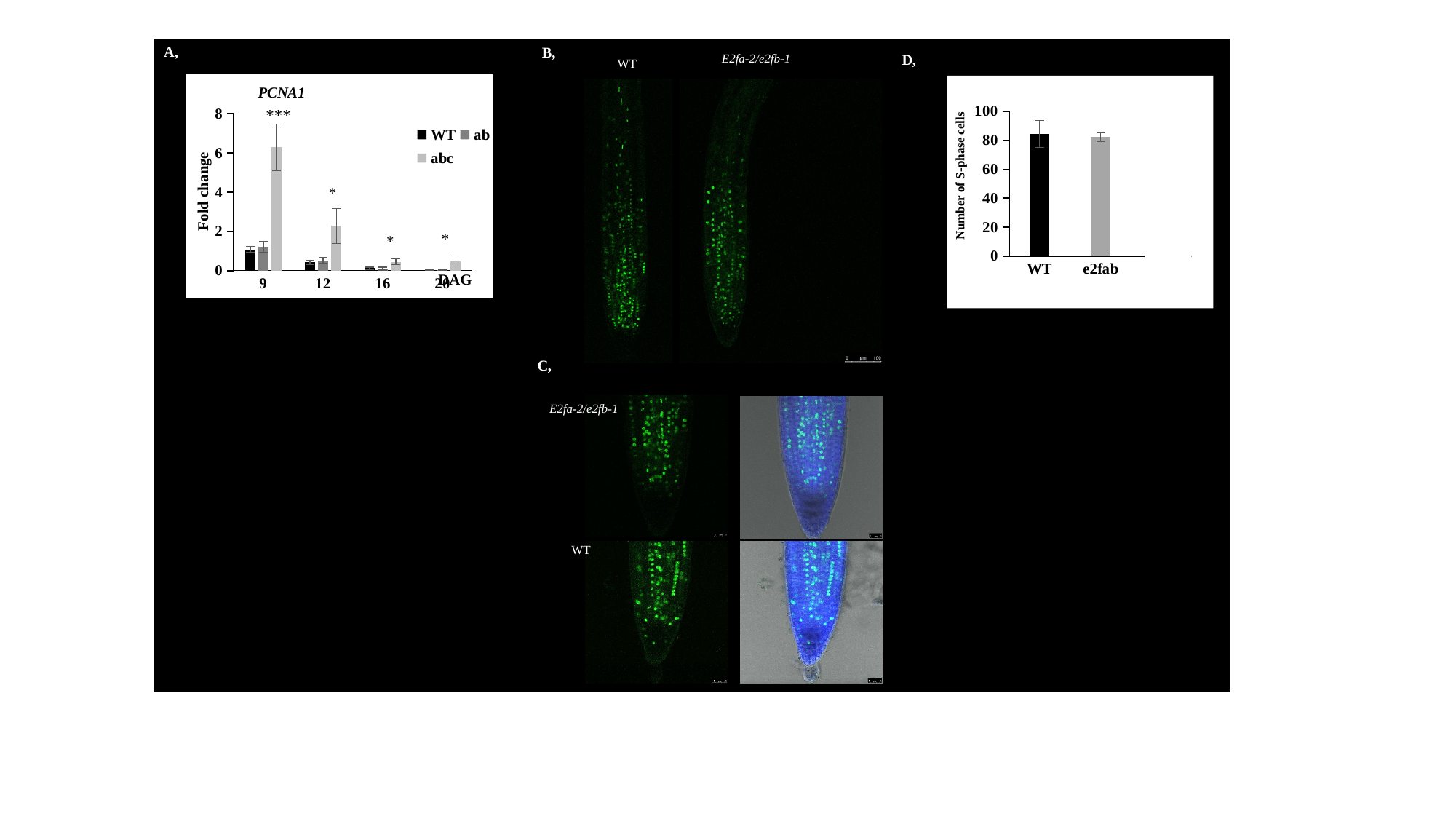

A,
B,
E2fa-2/e2fb-1
WT
D,
#### Chart: PCNA1
| Category | | | |
|---|---|---|---|
| 9 | 1.0736001532936923 | 1.2112092660241134 | 6.282730053357251 |
| 12 | 0.4226618686443239 | 0.4985233839530281 | 2.269078903579262 |
| 16 | 0.11895139617394722 | 0.08928155442678394 | 0.44623996294901586 |
| 20 | 0.036391418754562525 | 0.031743660964784766 | 0.4836523461805593 |***
Fold change
DAG
#### Chart
| Category | |
|---|---|
| WT | 84.3 |
| e2fab | 82.5 |
Number of S-phase cells
C,
E2fa-2/e2fb-1
WT

### Slide 3
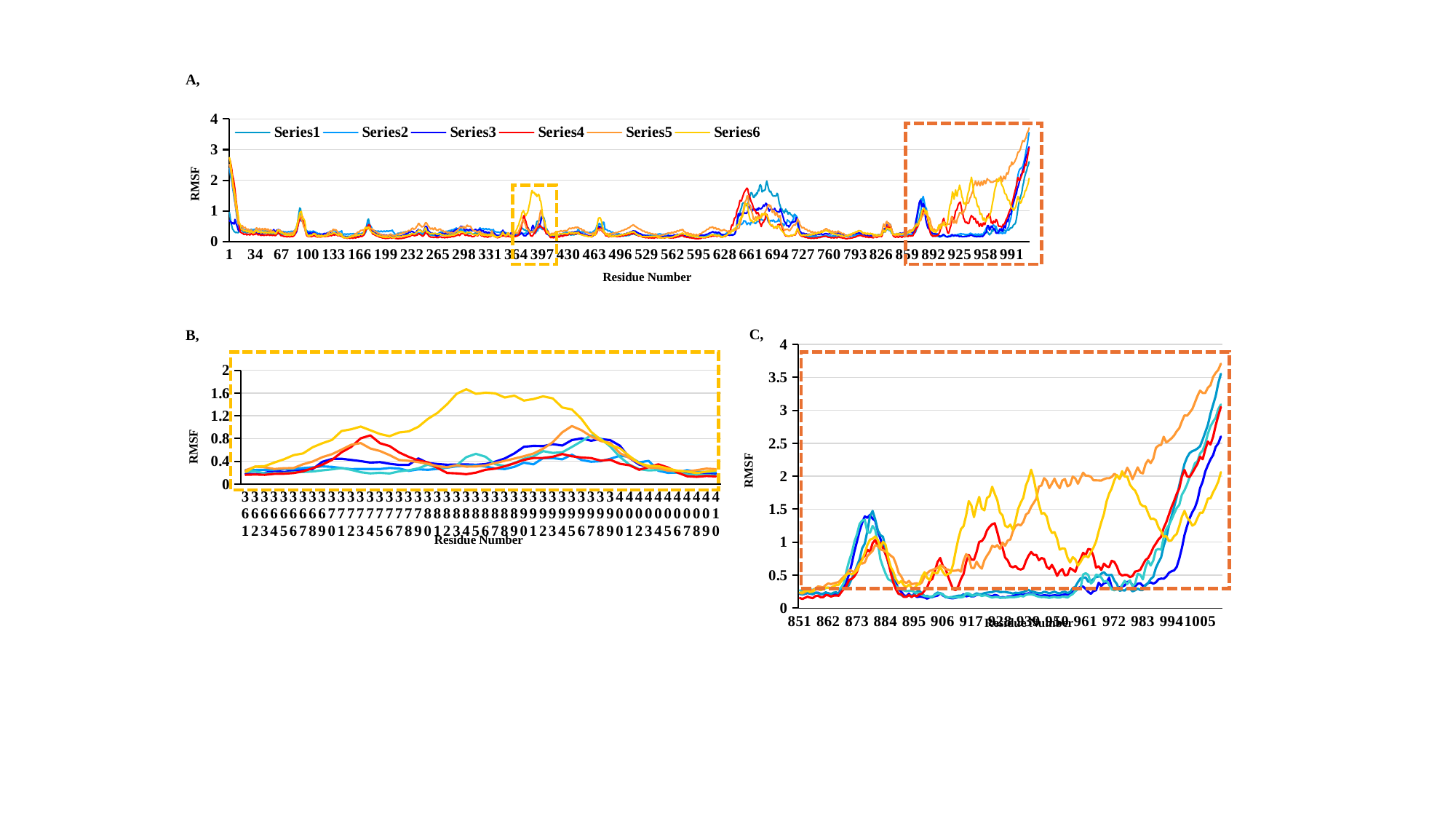

A,
#### Chart
| Category | Series1 | Series2 | Series3 | | Series4 | Series5 | Series6 |
|---|---|---|---|---|---|---|---|
RMSF
Residue Number
C,
B,
#### Chart
| Category | Series1 | Series2 | Series3 | | Series4 | Series5 | Series6 |
|---|---|---|---|---|---|---|---|
| 851 | 0.2166 | 0.2191 | 0.2697 | None | 0.1528 | 0.2293 | 0.2343 |
| 852 | 0.2057 | 0.2059 | 0.2585 | None | 0.1378 | 0.2214 | 0.2339 |
| 853 | 0.2161 | 0.219 | 0.2715 | None | 0.1557 | 0.2431 | 0.2599 |
| 854 | 0.2294 | 0.232 | 0.2913 | None | 0.1729 | 0.2668 | 0.2624 |
| 855 | 0.2177 | 0.2195 | 0.2808 | None | 0.1584 | 0.2694 | 0.2427 |
| 856 | 0.2113 | 0.2113 | 0.2768 | None | 0.151 | 0.2723 | 0.2514 |
| 857 | 0.2309 | 0.234 | 0.2959 | None | 0.1807 | 0.2935 | 0.2801 |
| 858 | 0.2336 | 0.2287 | 0.3122 | None | 0.188 | 0.3274 | 0.2859 |
| 859 | 0.2133 | 0.2069 | 0.2935 | None | 0.1645 | 0.3224 | 0.2719 |
| 860 | 0.2101 | 0.219 | 0.2837 | None | 0.1654 | 0.3121 | 0.2866 |
| 861 | 0.2271 | 0.2392 | 0.3027 | None | 0.1958 | 0.3497 | 0.3113 |
| 862 | 0.2242 | 0.2221 | 0.3108 | None | 0.1926 | 0.3734 | 0.3083 |
| 863 | 0.2103 | 0.2111 | 0.2949 | None | 0.1739 | 0.3624 | 0.308 |
| 864 | 0.2205 | 0.234 | 0.2998 | None | 0.188 | 0.3744 | 0.33 |
| 865 | 0.2213 | 0.2392 | 0.2896 | None | 0.1899 | 0.3858 | 0.3494 |
| 866 | 0.216 | 0.2181 | 0.2795 | None | 0.1874 | 0.3941 | 0.3541 |
| 867 | 0.3095 | 0.2656 | 0.3157 | None | 0.2452 | 0.4391 | 0.4063 |
| 868 | 0.3288 | 0.2957 | 0.3841 | None | 0.2889 | 0.4786 | 0.4532 |
| 869 | 0.4026 | 0.2921 | 0.5688 | None | 0.3393 | 0.5122 | 0.5098 |
| 870 | 0.5216 | 0.3605 | 0.7196 | None | 0.4279 | 0.5793 | 0.5258 |
| 871 | 0.6751 | 0.4487 | 0.8342 | None | 0.4428 | 0.5621 | 0.5109 |
| 872 | 0.8583 | 0.5326 | 1.0045 | None | 0.4819 | 0.5404 | 0.5292 |
| 873 | 1.0116 | 0.6398 | 1.1208 | None | 0.5487 | 0.6102 | 0.5698 |
| 874 | 1.1713 | 0.7405 | 1.274 | None | 0.6652 | 0.674 | 0.663 |
| 875 | 1.2957 | 0.9074 | 1.3264 | None | 0.7474 | 0.6744 | 0.7294 |
| 876 | 1.3903 | 0.9772 | 1.34 | None | 0.7935 | 0.6911 | 0.8361 |
| 877 | 1.3682 | 1.1688 | 1.1521 | None | 0.8851 | 0.7884 | 0.9596 |
| 878 | 1.4142 | 1.3925 | 1.1413 | None | 0.8503 | 0.8268 | 1.0373 |
| 879 | 1.3564 | 1.4731 | 1.245 | None | 0.9847 | 0.8664 | 1.0512 |
| 880 | 1.3245 | 1.3467 | 1.1885 | None | 1.0325 | 0.955 | 1.0859 |
| 881 | 1.1666 | 1.1788 | 0.9911 | None | 0.9396 | 0.9427 | 1.0328 |
| 882 | 1.0537 | 1.1061 | 0.7419 | None | 0.9896 | 0.8906 | 0.9213 |
| 883 | 0.929 | 1.081 | 0.6231 | None | 0.9102 | 0.8771 | 1.0067 |
| 884 | 0.8156 | 0.9 | 0.5236 | None | 0.8215 | 0.8561 | 0.9588 |
| 885 | 0.7399 | 0.6791 | 0.4379 | None | 0.7022 | 0.8286 | 0.7648 |
| 886 | 0.6004 | 0.6121 | 0.4184 | None | 0.534 | 0.7943 | 0.6229 |
| 887 | 0.4392 | 0.4356 | 0.3829 | None | 0.3842 | 0.7653 | 0.5497 |
| 888 | 0.3761 | 0.2972 | 0.3358 | None | 0.2812 | 0.6686 | 0.435 |
| 889 | 0.2885 | 0.2218 | 0.3063 | None | 0.2141 | 0.5342 | 0.3752 |
| 890 | 0.2492 | 0.1975 | 0.2866 | None | 0.2119 | 0.4825 | 0.4085 |
| 891 | 0.1948 | 0.1738 | 0.2669 | None | 0.1693 | 0.3999 | 0.3472 |
| 892 | 0.1812 | 0.174 | 0.2551 | None | 0.1701 | 0.3798 | 0.3166 |
| 893 | 0.2184 | 0.194 | 0.2618 | None | 0.1963 | 0.4117 | 0.357 |
| 894 | 0.1707 | 0.1892 | 0.2618 | None | 0.1685 | 0.362 | 0.308 |
| 895 | 0.2137 | 0.1932 | 0.244 | None | 0.1936 | 0.3694 | 0.2969 |
| 896 | 0.1677 | 0.247 | 0.2304 | None | 0.1804 | 0.377 | 0.3458 |
| 897 | 0.1732 | 0.2611 | 0.2302 | None | 0.2095 | 0.356 | 0.3704 |
| 898 | 0.1709 | 0.2232 | 0.195 | None | 0.2155 | 0.3918 | 0.4751 |
| 899 | 0.1601 | 0.1791 | 0.1832 | None | 0.2661 | 0.48 | 0.5443 |
| 900 | 0.1422 | 0.185 | 0.1772 | None | 0.3153 | 0.528 | 0.4581 |
| 901 | 0.1636 | 0.1688 | 0.1735 | None | 0.4154 | 0.5644 | 0.4229 |
| 902 | 0.1717 | 0.1643 | 0.1793 | None | 0.4402 | 0.5772 | 0.5402 |
| 903 | 0.1805 | 0.2108 | 0.1968 | None | 0.5686 | 0.5951 | 0.5319 |
| 904 | 0.1894 | 0.2358 | 0.2203 | None | 0.697 | 0.6083 | 0.529 |
| 905 | 0.2252 | 0.2262 | 0.2233 | None | 0.76 | 0.6494 | 0.6244 |
| 906 | 0.1921 | 0.2096 | 0.2018 | None | 0.6378 | 0.6344 | 0.5504 |
| 907 | 0.1719 | 0.1661 | 0.1772 | None | 0.6031 | 0.5891 | 0.4958 |
| 908 | 0.1593 | 0.1599 | 0.1607 | None | 0.4835 | 0.5847 | 0.5164 |
| 909 | 0.1505 | 0.1651 | 0.1617 | None | 0.3787 | 0.5636 | 0.5364 |
| 910 | 0.1486 | 0.1695 | 0.1554 | None | 0.2842 | 0.5658 | 0.6648 |
| 911 | 0.1564 | 0.1791 | 0.1614 | None | 0.2715 | 0.5674 | 0.8703 |
| 912 | 0.1634 | 0.1876 | 0.1637 | None | 0.3352 | 0.5797 | 1.0519 |
| 913 | 0.1739 | 0.1888 | 0.1618 | None | 0.4432 | 0.5518 | 1.1991 |
| 914 | 0.2122 | 0.2011 | 0.1714 | None | 0.5181 | 0.7175 | 1.2424 |
| 915 | 0.1771 | 0.2065 | 0.2244 | None | 0.7002 | 0.8182 | 1.4087 |
| 916 | 0.1897 | 0.1963 | 0.2226 | None | 0.8055 | 0.7748 | 1.6223 |
| 917 | 0.179 | 0.1764 | 0.199 | None | 0.7362 | 0.6116 | 1.5555 |
| 918 | 0.1783 | 0.2012 | 0.1872 | None | 0.7313 | 0.6043 | 1.3793 |
| 919 | 0.1994 | 0.2231 | 0.2031 | None | 0.8514 | 0.7036 | 1.5537 |
| 920 | 0.2051 | 0.2111 | 0.2019 | None | 1.0033 | 0.6354 | 1.6858 |
| 921 | 0.1956 | 0.2101 | 0.1834 | None | 1.0164 | 0.598 | 1.5108 |
| 922 | 0.2063 | 0.2263 | 0.1933 | None | 1.0694 | 0.724 | 1.4813 |
| 923 | 0.193 | 0.2337 | 0.1915 | None | 1.1778 | 0.7943 | 1.6698 |
| 924 | 0.189 | 0.2386 | 0.173 | None | 1.2334 | 0.8521 | 1.6978 |
| 925 | 0.1812 | 0.2411 | 0.1583 | None | 1.2716 | 0.9448 | 1.84 |
| 926 | 0.1995 | 0.2583 | 0.1684 | None | 1.2857 | 0.9246 | 1.7319 |
| 927 | 0.1906 | 0.2571 | 0.1687 | None | 1.1508 | 0.9548 | 1.6347 |
| 928 | 0.1566 | 0.2419 | 0.1582 | None | 0.9955 | 0.8977 | 1.4498 |
| 929 | 0.1673 | 0.2441 | 0.157 | None | 0.9154 | 0.9933 | 1.4051 |
| 930 | 0.1604 | 0.2447 | 0.1595 | None | 0.7663 | 0.946 | 1.2488 |
| 931 | 0.1719 | 0.2363 | 0.1598 | None | 0.7226 | 1.0255 | 1.225 |
| 932 | 0.1769 | 0.2321 | 0.1659 | None | 0.6354 | 1.0398 | 1.2663 |
| 933 | 0.1801 | 0.2204 | 0.1632 | None | 0.6183 | 1.1631 | 1.186 |
| 934 | 0.1965 | 0.2325 | 0.167 | None | 0.6377 | 1.2432 | 1.3263 |
| 935 | 0.1947 | 0.2281 | 0.1739 | None | 0.5949 | 1.2683 | 1.5021 |
| 936 | 0.2065 | 0.236 | 0.1862 | None | 0.5846 | 1.2538 | 1.5987 |
| 937 | 0.201 | 0.2396 | 0.1756 | None | 0.6012 | 1.3069 | 1.6746 |
| 938 | 0.2113 | 0.2616 | 0.1997 | None | 0.7114 | 1.4171 | 1.8599 |
| 939 | 0.2236 | 0.2744 | 0.2071 | None | 0.7924 | 1.4464 | 1.9559 |
| 940 | 0.2193 | 0.2529 | 0.2065 | None | 0.8509 | 1.5339 | 2.0997 |
| 941 | 0.2026 | 0.2457 | 0.2014 | None | 0.8062 | 1.5959 | 1.9637 |
| 942 | 0.2002 | 0.234 | 0.1843 | None | 0.8072 | 1.6587 | 1.7643 |
| 943 | 0.1902 | 0.2259 | 0.1717 | None | 0.7256 | 1.8425 | 1.5729 |
| 944 | 0.1876 | 0.2286 | 0.1657 | None | 0.7594 | 1.8595 | 1.4308 |
| 945 | 0.1944 | 0.246 | 0.168 | None | 0.7443 | 1.9717 | 1.4421 |
| 946 | 0.1913 | 0.2386 | 0.1623 | None | 0.6259 | 1.9302 | 1.3875 |
| 947 | 0.1877 | 0.2247 | 0.1549 | None | 0.5933 | 1.8206 | 1.2202 |
| 948 | 0.1895 | 0.2358 | 0.1646 | None | 0.655 | 1.8881 | 1.1386 |
| 949 | 0.1941 | 0.2478 | 0.1704 | None | 0.5837 | 1.9641 | 1.1534 |
| 950 | 0.1916 | 0.2316 | 0.1592 | None | 0.4884 | 1.8721 | 1.0511 |
| 951 | 0.1914 | 0.2218 | 0.1583 | None | 0.5557 | 1.8183 | 0.8858 |
| 952 | 0.2001 | 0.2416 | 0.1739 | None | 0.592 | 1.9359 | 0.9052 |
| 953 | 0.2071 | 0.2477 | 0.1688 | None | 0.4968 | 1.9577 | 0.9018 |
| 954 | 0.2028 | 0.229 | 0.1592 | None | 0.5033 | 1.8459 | 0.765 |
| 955 | 0.2215 | 0.2414 | 0.1902 | None | 0.6047 | 1.8693 | 0.6916 |
| 956 | 0.2438 | 0.3016 | 0.2121 | None | 0.5811 | 1.9944 | 0.7723 |
| 957 | 0.2599 | 0.3222 | 0.2571 | None | 0.5506 | 1.9674 | 0.7404 |
| 958 | 0.2993 | 0.3931 | 0.2833 | None | 0.6617 | 1.8847 | 0.6436 |
| 959 | 0.3213 | 0.4496 | 0.2976 | None | 0.7426 | 1.9713 | 0.6962 |
| 960 | 0.3266 | 0.4592 | 0.5084 | None | 0.8384 | 2.0537 | 0.7695 |
| 961 | 0.2893 | 0.4641 | 0.5276 | None | 0.8057 | 2.0088 | 0.7915 |
| 962 | 0.2448 | 0.3973 | 0.5016 | None | 0.897 | 2.0075 | 0.7696 |
| 963 | 0.2161 | 0.4115 | 0.3724 | None | 0.886 | 1.9917 | 0.8471 |
| 964 | 0.2593 | 0.4403 | 0.4089 | None | 0.7852 | 1.9376 | 0.9177 |
| 965 | 0.2702 | 0.4717 | 0.5072 | None | 0.6131 | 1.9386 | 1.0097 |
| 966 | 0.3845 | 0.4771 | 0.4952 | None | 0.6348 | 1.9365 | 1.1638 |
| 967 | 0.3346 | 0.5214 | 0.4633 | None | 0.5776 | 1.9343 | 1.3016 |
| 968 | 0.3753 | 0.5449 | 0.3961 | None | 0.6736 | 1.9556 | 1.4259 |
| 969 | 0.3883 | 0.5031 | 0.4114 | None | 0.6342 | 1.9694 | 1.614 |
| 970 | 0.4599 | 0.4987 | 0.3541 | None | 0.6208 | 1.9704 | 1.7342 |
| 971 | 0.3172 | 0.5047 | 0.2829 | None | 0.7177 | 1.9889 | 1.8149 |
| 972 | 0.29 | 0.4166 | 0.2701 | None | 0.7002 | 2.0378 | 1.9387 |
| 973 | 0.2852 | 0.3658 | 0.284 | None | 0.6298 | 2.0106 | 2.0181 |
| 974 | 0.3157 | 0.2624 | 0.2804 | None | 0.5226 | 1.9593 | 1.9626 |
| 975 | 0.3258 | 0.2721 | 0.3495 | None | 0.4944 | 2.035 | 2.0765 |
| 976 | 0.3434 | 0.266 | 0.412 | None | 0.5033 | 2.0305 | 1.9929 |
| 977 | 0.3973 | 0.2997 | 0.3647 | None | 0.5009 | 2.1297 | 1.9948 |
| 978 | 0.4052 | 0.2903 | 0.421 | None | 0.4693 | 2.0569 | 1.8731 |
| 979 | 0.3445 | 0.2538 | 0.3329 | None | 0.4856 | 1.9551 | 1.8185 |
| 980 | 0.3293 | 0.2692 | 0.3686 | None | 0.554 | 2.0436 | 1.781 |
| 981 | 0.3687 | 0.2952 | 0.5197 | None | 0.562 | 2.1349 | 1.6969 |
| 982 | 0.376 | 0.2737 | 0.5046 | None | 0.5814 | 2.0578 | 1.5803 |
| 983 | 0.3356 | 0.2729 | 0.434 | None | 0.6579 | 2.0429 | 1.5474 |
| 984 | 0.343 | 0.3494 | 0.6159 | None | 0.7288 | 2.1825 | 1.5415 |
| 985 | 0.3814 | 0.3774 | 0.7164 | None | 0.7587 | 2.2465 | 1.4421 |
| 986 | 0.3898 | 0.441 | 0.645 | None | 0.8285 | 2.1992 | 1.3517 |
| 987 | 0.3708 | 0.474 | 0.7212 | None | 0.9187 | 2.2679 | 1.3571 |
| 988 | 0.3901 | 0.6101 | 0.8803 | None | 0.9782 | 2.4303 | 1.3326 |
| 989 | 0.4358 | 0.6944 | 0.8958 | None | 1.0406 | 2.4664 | 1.2254 |
| 990 | 0.4472 | 0.7686 | 0.8853 | None | 1.0814 | 2.4735 | 1.1656 |
| 991 | 0.4449 | 0.9415 | 1.0685 | None | 1.2146 | 2.5996 | 1.0814 |
| 992 | 0.4778 | 1.0657 | 1.1977 | None | 1.3032 | 2.512 | 1.0912 |
| 993 | 0.5344 | 1.2466 | 1.2579 | None | 1.4159 | 2.5415 | 1.0178 |
| 994 | 0.5578 | 1.3912 | 1.3399 | None | 1.5299 | 2.5745 | 1.028 |
| 995 | 0.5709 | 1.5321 | 1.4302 | None | 1.6239 | 2.6185 | 1.0895 |
| 996 | 0.624 | 1.6805 | 1.517 | None | 1.732 | 2.6823 | 1.1242 |
| 997 | 0.751 | 1.8643 | 1.5654 | None | 1.8116 | 2.7289 | 1.2424 |
| 998 | 0.911 | 2.0238 | 1.72 | None | 1.997 | 2.8329 | 1.3876 |
| 999 | 1.1065 | 2.1871 | 1.7864 | None | 2.0955 | 2.9237 | 1.4733 |
| 1000 | 1.2457 | 2.2865 | 1.8769 | None | 1.9975 | 2.9185 | 1.3747 |
| 1001 | 1.3647 | 2.3525 | 1.991 | None | 1.9891 | 2.9648 | 1.3242 |
| 1002 | 1.4546 | 2.3813 | 2.0859 | None | 2.0512 | 3.0189 | 1.2504 |
| 1003 | 1.522 | 2.3954 | 2.1959 | None | 2.1215 | 3.1167 | 1.275 |
| 1004 | 1.6364 | 2.4208 | 2.2288 | None | 2.1897 | 3.2193 | 1.3666 |
| 1005 | 1.8175 | 2.4558 | 2.3445 | None | 2.2951 | 3.2988 | 1.4469 |
| 1006 | 1.9188 | 2.5572 | 2.3934 | None | 2.2573 | 3.2629 | 1.4464 |
| 1007 | 2.0727 | 2.6672 | 2.4949 | None | 2.4059 | 3.2621 | 1.5391 |
| 1008 | 2.1686 | 2.7804 | 2.6458 | None | 2.5262 | 3.3434 | 1.6604 |
| 1009 | 2.2553 | 2.9493 | 2.7571 | None | 2.4805 | 3.3804 | 1.6635 |
| 1010 | 2.3205 | 3.085 | 2.8282 | None | 2.5985 | 3.5063 | 1.7563 |
| 1011 | 2.4443 | 3.2158 | 2.8886 | None | 2.7697 | 3.5674 | 1.8283 |
| 1012 | 2.4977 | 3.4116 | 3.0178 | None | 2.9187 | 3.6135 | 1.9195 |
| 1013 | 2.6026 | 3.5538 | 3.0884 | None | 3.0514 | 3.7066 | 2.062 |
RMSF
Residue Number
#### Chart
| Category | Series1 | Series2 | Series3 | | Series4 | Series5 | Series6 |
|---|---|---|---|---|---|---|---|
| 361 | 0.18 | 0.2467 | 0.2144 | None | 0.1611 | 0.2396 | 0.2118 |
| 362 | 0.173 | 0.2464 | 0.2207 | None | 0.1666 | 0.3056 | 0.3051 |
| 363 | 0.1957 | 0.2498 | 0.1884 | None | 0.1616 | 0.2946 | 0.3143 |
| 364 | 0.2327 | 0.2514 | 0.1874 | None | 0.178 | 0.2633 | 0.3767 |
| 365 | 0.2108 | 0.2646 | 0.1925 | None | 0.1805 | 0.277 | 0.4345 |
| 366 | 0.2319 | 0.2633 | 0.2042 | None | 0.1927 | 0.2803 | 0.5068 |
| 367 | 0.2526 | 0.2802 | 0.2116 | None | 0.2236 | 0.3474 | 0.538 |
| 368 | 0.2627 | 0.2921 | 0.2197 | None | 0.2779 | 0.395 | 0.6475 |
| 369 | 0.3907 | 0.3094 | 0.2403 | None | 0.34 | 0.4723 | 0.7169 |
| 370 | 0.44 | 0.3014 | 0.2573 | None | 0.4232 | 0.5243 | 0.7752 |
| 371 | 0.4433 | 0.2793 | 0.2798 | None | 0.5577 | 0.6053 | 0.9322 |
| 372 | 0.4229 | 0.2619 | 0.2493 | None | 0.6518 | 0.69 | 0.9629 |
| 373 | 0.4019 | 0.2632 | 0.2071 | None | 0.8044 | 0.7199 | 1.0109 |
| 374 | 0.3759 | 0.2633 | 0.1862 | None | 0.8558 | 0.6248 | 0.9453 |
| 375 | 0.3846 | 0.2625 | 0.1971 | None | 0.7177 | 0.5805 | 0.8795 |
| 376 | 0.356 | 0.2817 | 0.1846 | None | 0.6679 | 0.5079 | 0.841 |
| 377 | 0.3352 | 0.2719 | 0.2217 | None | 0.5544 | 0.4216 | 0.9072 |
| 378 | 0.3362 | 0.231 | 0.2417 | None | 0.4768 | 0.4077 | 0.9256 |
| 379 | 0.4501 | 0.2571 | 0.2747 | None | 0.4133 | 0.3845 | 1.0068 |
| 380 | 0.3687 | 0.2502 | 0.3416 | None | 0.3783 | 0.3545 | 1.1462 |
| 381 | 0.3515 | 0.2654 | 0.2773 | None | 0.2813 | 0.3141 | 1.252 |
| 382 | 0.3386 | 0.2873 | 0.2974 | None | 0.1937 | 0.2922 | 1.4043 |
| 383 | 0.3421 | 0.3118 | 0.3286 | None | 0.1862 | 0.3293 | 1.5878 |
| 384 | 0.3447 | 0.3203 | 0.4721 | None | 0.1723 | 0.3051 | 1.6674 |
| 385 | 0.3375 | 0.3304 | 0.5303 | None | 0.1996 | 0.3081 | 1.5865 |
| 386 | 0.3501 | 0.3067 | 0.4791 | None | 0.25 | 0.3274 | 1.6062 |
| 387 | 0.3914 | 0.2744 | 0.3483 | None | 0.2699 | 0.3675 | 1.5954 |
| 388 | 0.4488 | 0.2618 | 0.3155 | None | 0.3107 | 0.4066 | 1.522 |
| 389 | 0.5376 | 0.3015 | 0.3671 | None | 0.364 | 0.4464 | 1.5539 |
| 390 | 0.6541 | 0.3747 | 0.4439 | None | 0.4239 | 0.488 | 1.4675 |
| 391 | 0.6711 | 0.3458 | 0.487 | None | 0.4572 | 0.5355 | 1.4954 |
| 392 | 0.6666 | 0.4559 | 0.5778 | None | 0.4568 | 0.6206 | 1.5424 |
| 393 | 0.7012 | 0.4553 | 0.546 | None | 0.4788 | 0.7355 | 1.5064 |
| 394 | 0.6771 | 0.4334 | 0.5575 | None | 0.5265 | 0.9103 | 1.3454 |
| 395 | 0.7733 | 0.5156 | 0.6524 | None | 0.487 | 1.02 | 1.3117 |
| 396 | 0.8028 | 0.4213 | 0.7514 | None | 0.4657 | 0.9466 | 1.1431 |
| 397 | 0.7616 | 0.3915 | 0.8566 | None | 0.4558 | 0.8368 | 0.9186 |
| 398 | 0.7923 | 0.401 | 0.7992 | None | 0.4107 | 0.7562 | 0.7808 |
| 399 | 0.771 | 0.4428 | 0.6579 | None | 0.425 | 0.7209 | 0.6852 |
| 400 | 0.6741 | 0.497 | 0.475 | None | 0.3551 | 0.5423 | 0.6277 |
| 401 | 0.4578 | 0.4761 | 0.3394 | None | 0.329 | 0.4557 | 0.4905 |
| 402 | 0.3415 | 0.3796 | 0.2619 | None | 0.2497 | 0.3669 | 0.3666 |
| 403 | 0.2971 | 0.4067 | 0.2383 | None | 0.2971 | 0.2908 | 0.3197 |
| 404 | 0.2912 | 0.2326 | 0.2468 | None | 0.346 | 0.2802 | 0.3131 |
| 405 | 0.2363 | 0.1962 | 0.2455 | None | 0.2902 | 0.235 | 0.2729 |
| 406 | 0.204 | 0.2003 | 0.1958 | None | 0.2004 | 0.2342 | 0.2327 |
| 407 | 0.1795 | 0.2429 | 0.1667 | None | 0.1357 | 0.2192 | 0.2161 |
| 408 | 0.1774 | 0.2135 | 0.1616 | None | 0.1256 | 0.2434 | 0.1915 |
| 409 | 0.1834 | 0.2109 | 0.1588 | None | 0.14 | 0.2706 | 0.2175 |
| 410 | 0.1811 | 0.2045 | 0.1478 | None | 0.1338 | 0.2605 | 0.2346 |RMSF
Residue Number

### Slide 4
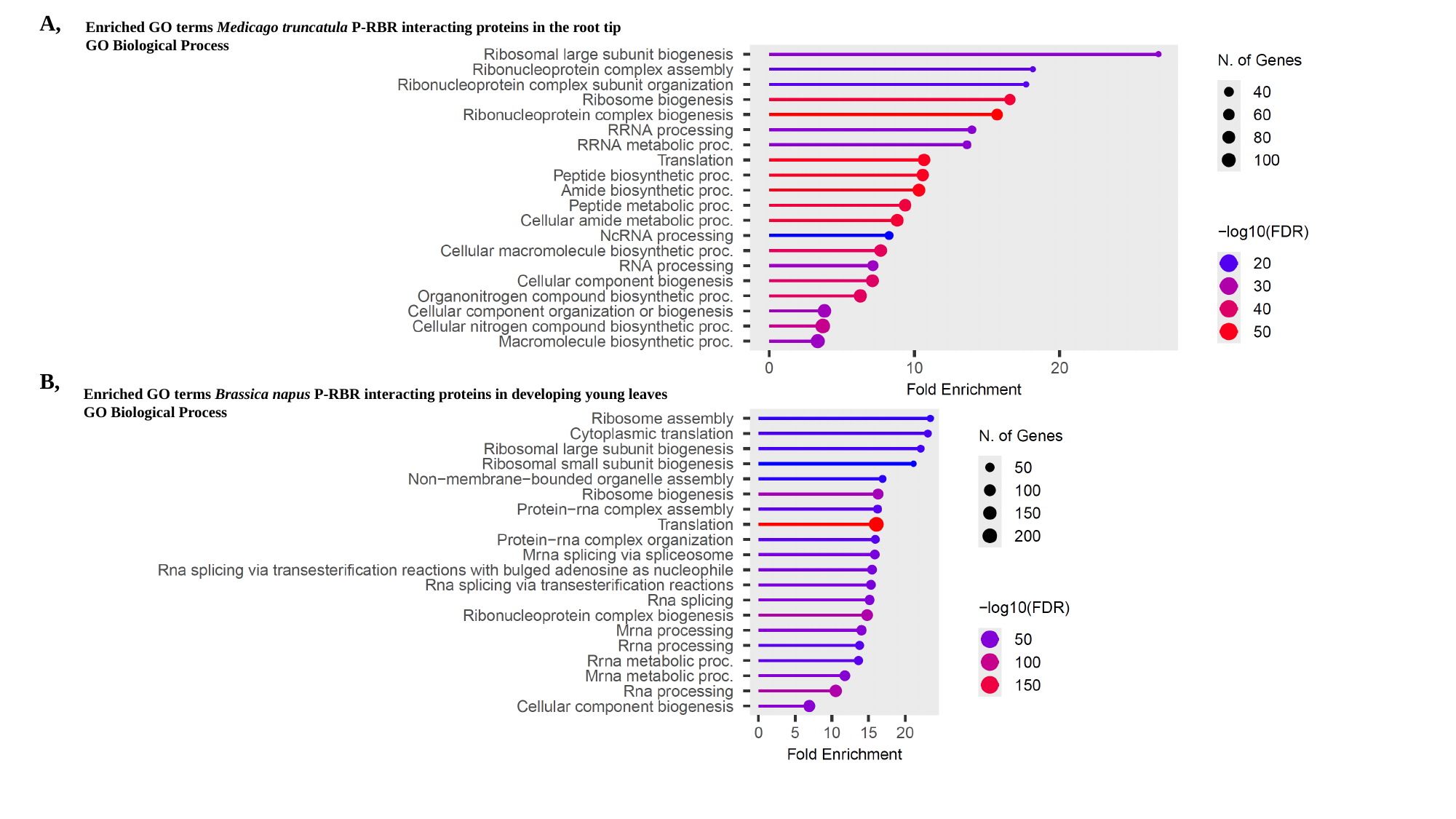

A,
Enriched GO terms Medicago truncatula P-RBR interacting proteins in the root tip
GO Biological Process
B,
Enriched GO terms Brassica napus P-RBR interacting proteins in developing young leaves
GO Biological Process

### Slide 5
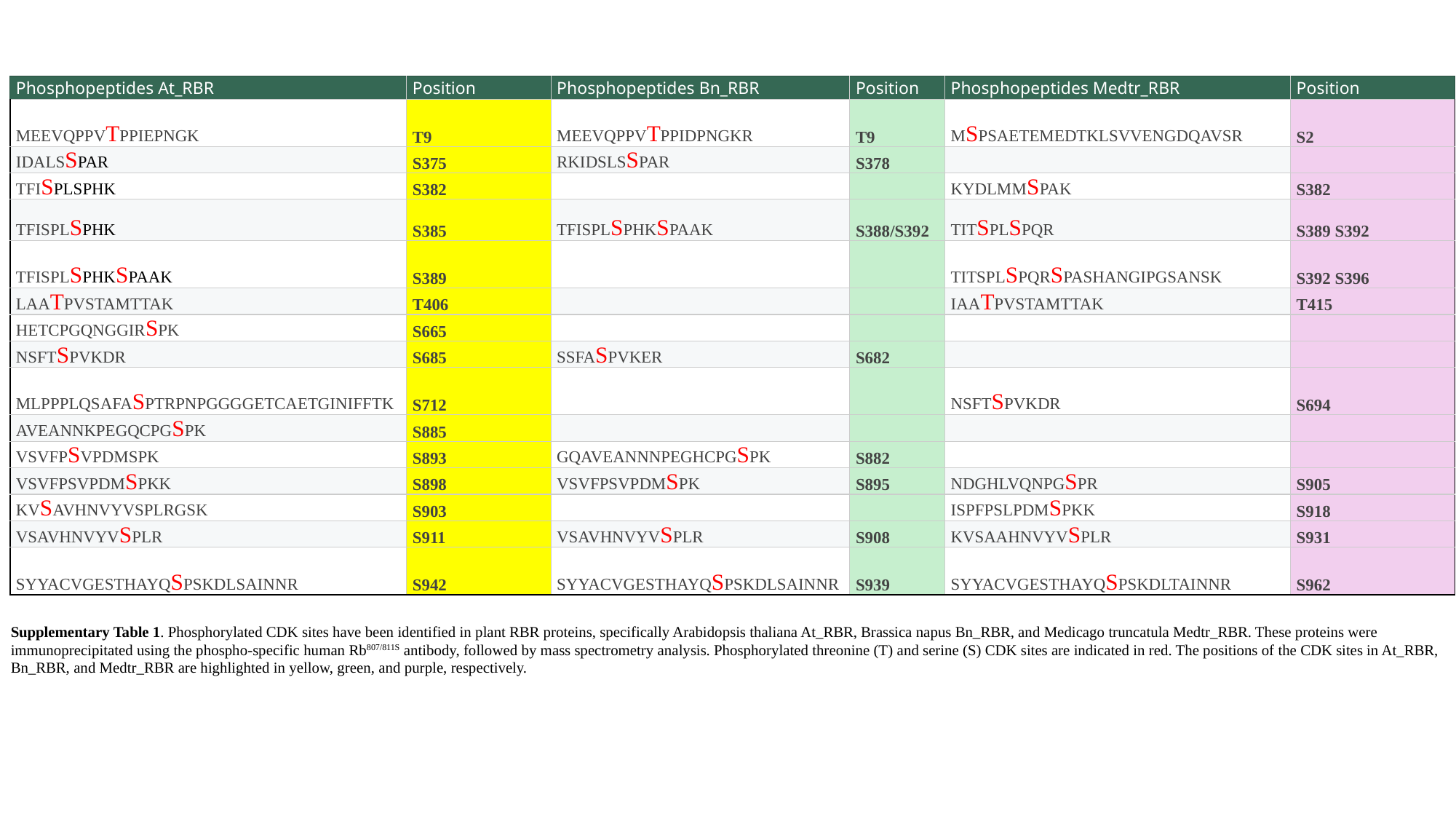

| Phosphopeptides At\_RBR | Position | Phosphopeptides Bn\_RBR | Position | Phosphopeptides Medtr\_RBR | Position |
| --- | --- | --- | --- | --- | --- |
| MEEVQPPVTPPIEPNGK | T9 | MEEVQPPVTPPIDPNGKR | T9 | MSPSAETEMEDTKLSVVENGDQAVSR | S2 |
| IDALSSPAR | S375 | RKIDSLSSPAR | S378 | | |
| TFISPLSPHK | S382 | | | KYDLMMSPAK | S382 |
| TFISPLSPHK | S385 | TFISPLSPHKSPAAK | S388/S392 | TITSPLSPQR | S389 S392 |
| TFISPLSPHKSPAAK | S389 | | | TITSPLSPQRSPASHANGIPGSANSK | S392 S396 |
| LAATPVSTAMTTAK | T406 | | | IAATPVSTAMTTAK | T415 |
| HETCPGQNGGIRSPK | S665 | | | | |
| NSFTSPVKDR | S685 | SSFASPVKER | S682 | | |
| MLPPPLQSAFASPTRPNPGGGGETCAETGINIFFTK | S712 | | | NSFTSPVKDR | S694 |
| AVEANNKPEGQCPGSPK | S885 | | | | |
| VSVFPSVPDMSPK | S893 | GQAVEANNNPEGHCPGSPK | S882 | | |
| VSVFPSVPDMSPKK | S898 | VSVFPSVPDMSPK | S895 | NDGHLVQNPGSPR | S905 |
| KVSAVHNVYVSPLRGSK | S903 | | | ISPFPSLPDMSPKK | S918 |
| VSAVHNVYVSPLR | S911 | VSAVHNVYVSPLR | S908 | KVSAAHNVYVSPLR | S931 |
| SYYACVGESTHAYQSPSKDLSAINNR | S942 | SYYACVGESTHAYQSPSKDLSAINNR | S939 | SYYACVGESTHAYQSPSKDLTAINNR | S962 |
Supplementary Table 1. Phosphorylated CDK sites have been identified in plant RBR proteins, specifically Arabidopsis thaliana At_RBR, Brassica napus Bn_RBR, and Medicago truncatula Medtr_RBR. These proteins were immunoprecipitated using the phospho-specific human Rb^807/811S^ antibody, followed by mass spectrometry analysis. Phosphorylated threonine (T) and serine (S) CDK sites are indicated in red. The positions of the CDK sites in At_RBR, Bn_RBR, and Medtr_RBR are highlighted in yellow, green, and purple, respectively.

### Slide 6
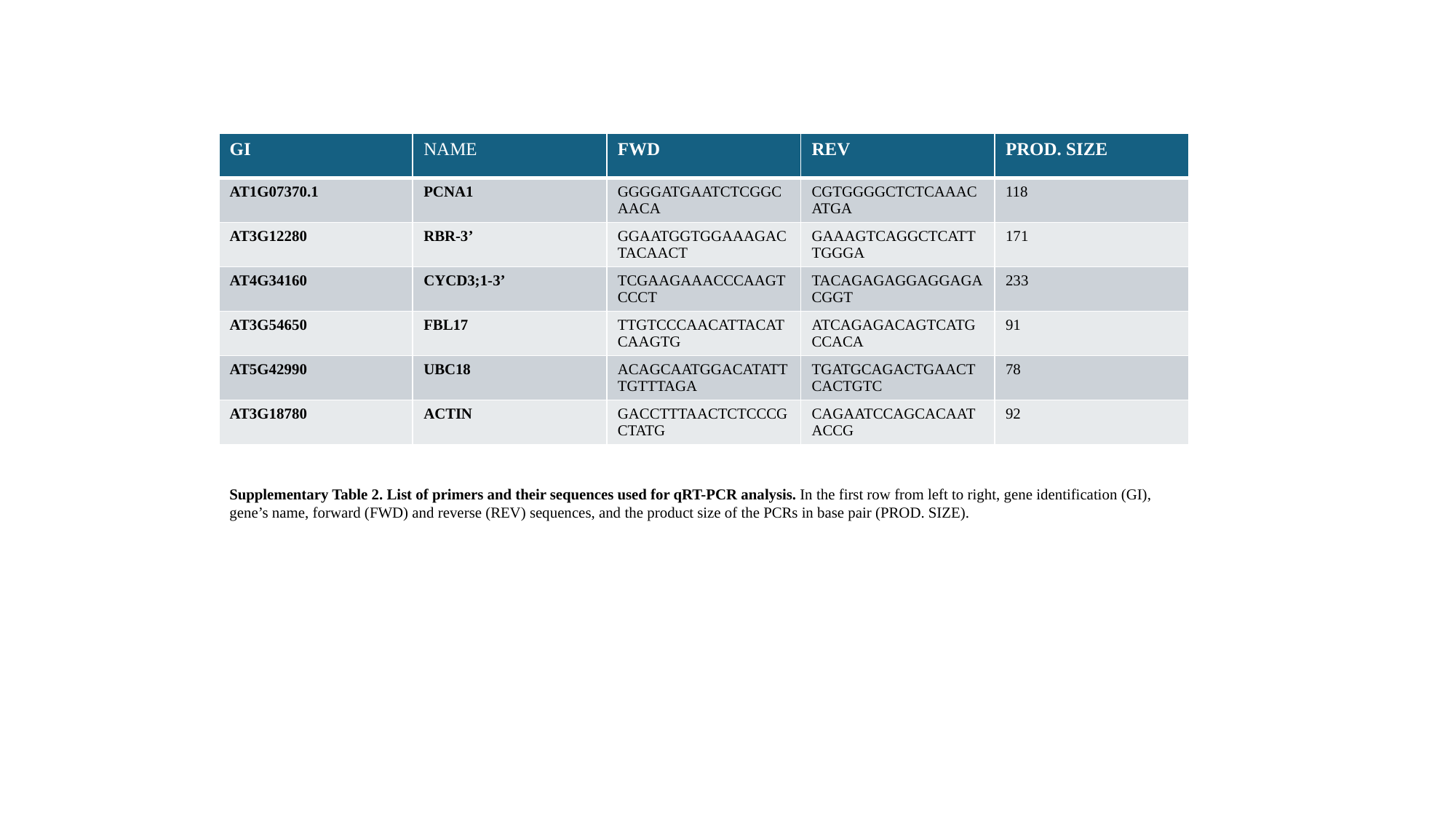

| GI | NAME | FWD | REV | PROD. SIZE |
| --- | --- | --- | --- | --- |
| AT1G07370.1 | PCNA1 | GGGGATGAATCTCGGCAACA | CGTGGGGCTCTCAAACATGA | 118 |
| AT3G12280 | RBR-3’ | GGAATGGTGGAAAGACTACAACT | GAAAGTCAGGCTCATTTGGGA | 171 |
| AT4G34160 | CYCD3;1-3’ | TCGAAGAAACCCAAGTCCCT | TACAGAGAGGAGGAGACGGT | 233 |
| AT3G54650 | FBL17 | TTGTCCCAACATTACATCAAGTG | ATCAGAGACAGTCATGCCACA | 91 |
| AT5G42990 | UBC18 | ACAGCAATGGACATATTTGTTTAGA | TGATGCAGACTGAACTCACTGTC | 78 |
| AT3G18780 | ACTIN | GACCTTTAACTCTCCCGCTATG | CAGAATCCAGCACAATACCG | 92 |
Supplementary Table 2. List of primers and their sequences used for qRT-PCR analysis. In the first row from left to right, gene identification (GI), gene’s name, forward (FWD) and reverse (REV) sequences, and the product size of the PCRs in base pair (PROD. SIZE).
